# Inhibitory Gain and Hub Architecture Confer Dynamic Resilience to Microcircuit Degeneration

**DOI:** 10.64898/2026.06.15.732346

**Authors:** Simachew Abebe Mengiste, Ad Aertsen, Arvind Kummar, Demian Battaglia

**Affiliations:** University of Strasbourg, Strasbourg, France; University of Freiburg, Freiburg im Breisgau, Germany; Royal Institute of Stockholm, Stockholm, Sweden

## Abstract

Neurodegeneration progressively removes synapses and neurons, yet neural circuits can retain stable collective dynamics despite substantial structural loss. Which structural principles confer this resilience remained unclear. Using large-scale spiking networks spanning empirical and synthetic microcircuit architectures, we systematically compared synaptic and neuronal modes of degeneration under controlled pruning strategies.

We found that resilience was not determined by connectivity loss alone, but by how inhibitory gain was embedded within circuit architecture. Networks in which inhibitory neurons occupied structurally central positions robustly maintained health-like firing rates, levels of synchrony, and informational bandwidth across degeneration stages, whereas architectures lacking such embedding exhibited amplified dynamical disruption.

Across regimes, the evolution of activity was organized by a compact set of weight-aware structural descriptors that generalized across network sizes and classes, with total effective synaptic coupling providing a dominant organizing axis. These results identified inhibitory architecture as a mechanistic determinant of circuit resilience and provided a predictive framework linking structural degeneration to collective dynamics.

## 1 Introduction

Neural circuits must maintain functional stability despite continuous structural changes. Across development and adulthood, synapses are formed, eliminated, and remodeled, and neuronal populations undergo turnover and reorganization. In brain diseases especially in neurodegenerative diseases, these processes are much amplified into progressive structural changes. The need for functional resilience raises the question of how much structural damage a network can sustain while preserving coherent activity dynamics. It is plausible that certain network architectures and the dynamical regimes they support may have evolved to confer greater resilience to degeneration.

Across neurodegenerative diseases, structural damage follows certain patterns. For instance, synaptic dysfunction and loss often precede overt neuronal death, initially leading to altered excitation–inhibition balance and hyperexcitability, followed by hypoactivity and circuit disconnection [1, 2, 3]. Selective vulnerability of projection neurons, dendritic spine loss, and remodeling of convergent synaptic inputs onto individual neurons have been documented in Alzheimer’s disease, Parkinson’s disease, Huntington’s disease, and amyotrophic lateral sclerosis (ALS) [4, 5, 6, 7]. Degeneration therefore reflects not simply a uniform reduction of connectivity but a redistribution of connectivity across specific synaptic and neuronal subpopulations.

At the network level, graph-theoretic analyses show that architectural features such as hub structure, degree heterogeneity, and modular organization influence robustness to damage [8, 9]. Preferential removal of hubs, peripheral nodes, or disruption of convergent synaptic inputs onto target neurons can lead to distinct structural outcomes even for similar levels of average connectivity loss. Yet network structure alone does not uniquely determine functional consequences. Networks with different wiring patterns can generate comparable activity regimes through homeostatic regulation and dynamical compensation [10, 11]. Functional resilience therefore depends not only on topology but on how connectivity shapes collective dynamics.

Cortical microcircuits typically operate in inhibition-dominated regimes characterized by irregular and largely asynchronous spiking despite strong recurrent coupling [12, 13, 14]. In these regimes, inhibitory gain control stabilizes excitation and constrains population variability. Crucially, the impact of inhibition depends on its architectural organization: the position of inhibitory neurons within the connectivity graph, their degree distribution, and the excitatory input they receive. These features determine how perturbations propagate through the circuit.

Understanding resilience under degeneration therefore requires integrating three elements: (i) structured patterns of synaptic and neuronal loss, (ii) the architecture of excitatory–inhibitory connectivity, and (iii) the dynamical regime governing population activity. Yet few studies have systematically compared degeneration modes across multiple network architectures while linking structural perturbations to multivariate activity outcomes through mechanistically interpretable descriptors.

Here we investigated how progressive degeneration reshaped activity dynamics in large-scale spiking networks and identified structural principles that conferred functional resilience. We compared simulations of an empirically reconstructed cortical microcircuit [15] with different models of complex random networks such as Erdos-Renyi random, small-world, and scale-free architectures. This approach allowed us to test whether naturally evolved connectivity exhibited nontrivial resilience relative to canonical models. We then applied controlled synaptic and neuronal pruning strategies capturing distinct modes of structural damage. All networks were initialized in comparable inhibition-dominated regimes, enabling controlled comparisons of degeneration responses.

We tracked how activity statistics evolved across degeneration stages, including firing rate, synchronization, and firing irregularity, a necessary condition for high informational bandwidth. We found that the empirical connectome was more resilient and maintained stable activity across a wide range of degeneration stages than most canonical-model architectures. This robustness was strongly associated with structurally central inhibitory hubs and elevated inhibitory gain. Beyond describing degeneration effects, we identified a compact set of weight-aware structural descriptors that predicted degeneration outcomes across architectures and pruning strategies.

These results provided a mechanistic account of circuit resilience in empirically reconstructed microcircuits. Functional stability under degeneration emerged not from network structure alone, but from its interaction with inhibitory gain control and the architectural embedding of excitatory–inhibitory connectivity.

## 2 Results

### 2.1 Network classes and baseline operating point

We considered six directed network classes with identical size and excitatory–inhibitory composition (80% excitatory, 20% inhibitory), operating in an inhibition-dominated regime. The reference network was based on an empirically reconstructed cortical microconnectome (`emp`; ***N* = 3611**) in which connections were estimated from three-dimensional soma distributions and axo-dendritic overlap statistics in layer 4 barrel cortex [15]. We compared this empirical reference with network classes featuring different random architectures: homogeneous Erdős–R’enyi networks (`er`), a smallworld topology (`sw`), and a scale-free class (`sf`). In addition, we considered modified versions of these networks in which inhibitory connectivity was strengthened to mimic features observed in empirical connectomes, as discussed later: an Erdős–R’enyi class preserving empirical excitatory–inhibitory subpopulation densities (`er-I`) and a scale-free class with an increased number of inhibitory hubs (`sf-I`). All classes preserved the total synapse count of the empirical reference but differed in degree heterogeneity and mesoscale organization of excitatory and inhibitory subpopulations.

Figure 1 summarized the structural organization and baseline dynamical characteristics of these classes. At the structural level (Figure 1A), the networks exhibited clear architectural differences. The empirical and scale-free classes displayed pronounced hub structure and heterogeneous connectivity, whereas Erdős–R’enyi and small-world networks were comparatively homogeneous. The subpopulation-constrained Erdős–R’enyi construction (`er-I`) preserved empirical inter-population densities but lacked the higher-order organization present in the empirical topology. In both the empirical network and the inhibition-structured variants, inhibitory neurons also occupied more central positions within the connectivity pattern, a feature that became important for interpreting degeneration responses in later sections.

**Figure 1.**
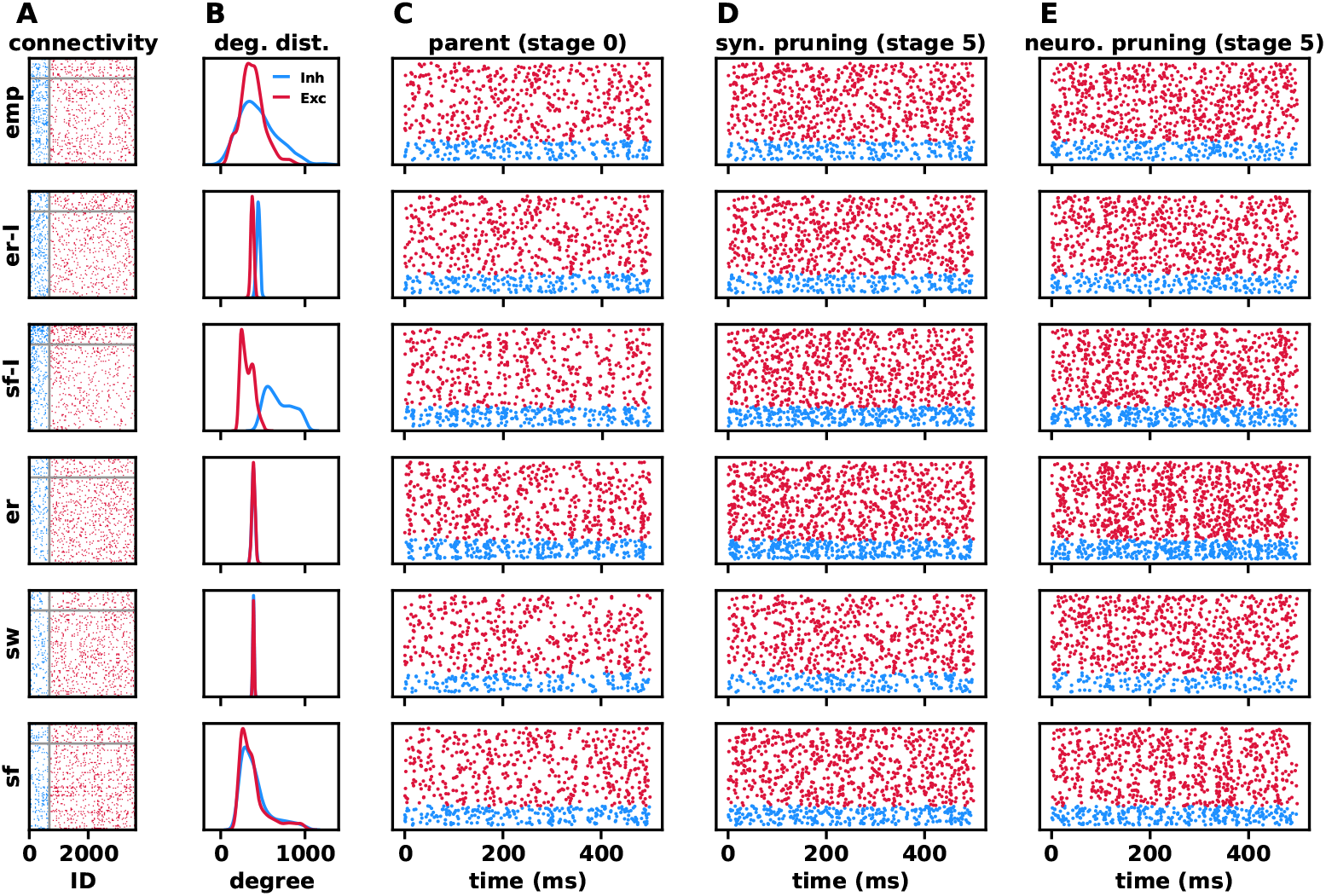
Network classes differ in structural organization while exhibiting comparable baseline activity. Six network classes are shown: empirical connectivity (`emp`), Erdős–Rényi with boosted inhibition architecture (`er-I`), scale-free with boosted inhibition architecture (`sf-I`), Erdős–Rényi (`er`), small-world (`sw`), and scale-free `(sf`) (rows). **(A)** Signed adjacency matrices illustrating architectural organization (columns denote sources, rows denote targets; red = excitatory, blue = inhibitory). **(B)** In-degree distributions for excitatory and inhibitory populations, highlighting differences in degree heterogeneity across classes. For representative degeneration-induced changes in these distributions, see Supplementary Fig. S1. **(C)** Representative raster plots of network activity in the intact baseline state (stage 0). **(D-E)** Example raster plots at intermediate degeneration (stage 5) under random synaptic and neuronal pruning (See next section). All networks were calibrated to comparable baseline firing rates, variability, and asynchrony prior to degeneration, allowing differences in degeneration trajectories to be attributed to architectural organization rather than baseline dynamical regimes.

These structural differences were reflected in the degree distributions (Figure 1B). Empirical and scale-free networks exhibited broad, heavy-tailed distributions for both excitatory and inhibitory populations, indicating substantial heterogeneity. In contrast, Erdős–R’enyi and small-world classes showed narrower distributions, consistent with more homogeneous connectivity. Importantly, the inhibition-structured scale-free class (`sf-I`) showed a pronounced broadening and rightward shift of the inhibitory degree distribution relative to the excitatory population, consistent with preferential embedding of inhibitory hubs. Thus, although `er-I` reproduced first-order subpopulation statistics, it did not capture the degree variability characteristic of the empirical network, whereas `sf-I` captured a stronger inhibitory-centrality signature.

To assess how these structural differences evolved under degeneration, we analyzed the corresponding degree distributions across pruning strategies and degeneration stages (Supplementary Fig. S1). While baseline differences reflected intrinsic architectural organization (Figure 1B), degeneration induced strategy-dependent redistributions of connectivity. Synaptic pruning primarily shifted and reshaped degree distributions while preserving population size, whereas neuronal pruning led to stronger distortions through node removal, including truncation of high-degree tails under hub-targeted strategies and narrowing under peripheral pruning.

Despite these structural differences, all networks were calibrated to operate in a comparable dynamical regime prior to degeneration (Figure 1C). In the intact state (stage 0), all classes exhibited asynchronous irregular activity with comparable firing rates, variability, and synchrony levels. In panel C, readers should note the similarly sparse rasters across all six rows, the irregular timing of spikes, and the absence of obvious population bursts or strong synchronous bands. This alignment ensured that subsequent differences observed under degeneration reflected intrinsic sensitivity to structural perturbations rather than baseline mismatches.

Under degeneration, network activity already diverged across network classes in the stage-5 examples shown in Figure 1D,E. Even at this intermediate stage under random pruning, structural differences translated into visibly distinct activity patterns, with some networks maintaining irregular firing while others showed clearer departures from the calibrated baseline. A systematic comparison of synaptic and neuronal pruning effects across strategies is presented in the following sections.

### 2.2 Degeneration strategies

To model the effects of neurodegeneration, we defined a set of procedures that progressively modified the initial (stage 0) networks through synapse ablation (*synaptic pruning* strategies) — emulating widespread synaptic dysfunction observed from early pathological stages [1, 2, 3, 16], often directly linked to functional impairment [17] — or through the removal of neurons (*neuronal death* strategies). We considered abstract degeneration mechanisms motivated by neurobiology without aiming to model any specific disease. The strategies spanned neutral, hub-targeted, peripheral-targeted, and broadcasting-targeted modes of structural damage; nevertheless, each pruning strategy had a possible neurophysiological interpretation.

We first examined five synaptic pruning strategies. In *random synaptic pruning* (s.rand), which served as a neutral reference, synapses were removed non-selectively. In *axonal pruning* (s.axon), we preferentially removed outgoing synapses from highly broadcasting neurons, emulating failure of strong projection outputs, as observed in amyotrophic lateral sclerosis (ALS), where early cortical hyperexcitability and disruption of inhibitory control were associated with degeneration of projection-heavy neurons [7, 18, 19]. *Dendritic pruning* (s.dend) instead targeted edges incident to highly innervated neurons, consistent with dendritic spine loss and remodeling of corticostriatal inputs reported in Parkinson’s disease (PD) and Huntington’s disease (HD) [20, 6, 21]. Conversely, *peripheral synaptic pruning* (`s.peri`) removed edges incident to weakly connected nodes, leading to early isolation of peripheral elements. Here and throughout, peripheral and central refer to topological roles in the network, with peripheral nodes corresponding to weakly connected (low-degree) nodes and central nodes to highly connected hubs, rather than spatial location or strength. This reflected synapse-first degeneration scenarios, particularly in Alzheimer’s disease, where subtle disconnection could precede overt neuronal loss [22]. Finally, *central synaptic pruning* (`s.cent`) targeted edges incident to hub neurons, leading to a more pronounced reduction in degree heterogeneity by preferentially truncating the high-degree tail of the distribution (see Supplementary Fig. S1). This served as a proxy for hub vulnerability and early inhibitory dysfunction, mechanisms implicated, for example, in cortical hyperexcitability and network instability in ALS [7, 19].

For the neuronal death scenarios, we adapted the synaptic pruning strategies for neurons. We again considered a neutral case of *random neuronal pruning* (`n.rand`), representing non-selective neuronal loss. We then introduced schemes in which neurons were preferentially removed according to either their overall connectivity (inputs and outputs) or, more specifically, their signal broadcasting capacity (outputs only). Thus, the distinction between central/peripheral and axonally central/axonally peripheral pruning was whether targeting was based on total degree or only on out-degree. *Peripheral neuronal pruning* (`n.peri`) and *central neuronal pruning* (`n.cent`) preferentially targeted nodes with, respectively, low and high numbers of connections. These corresponded to alternative hypotheses of hub resilience [2, 22] or hub fragility [5, 18] proposed for different neurodegenerative conditions. Finally, *axonally peripheral pruning* (`n.ax.peri`) and *axonally central pruning* (`n.ax.cent`) targeted weak and strong broadcasters, respectively, with the latter mimicking the vulnerability of projection-heavy neurons [7, 19].

All these processes were progressively applied to the different stage-0 networks to emulate the progression of neurodegeneration. Figure 1D-E provides representative stage-5 raster examples for the neutral synaptic and neuronal pruning schemes; the full effects of these pruning rules are quantified across classes and strategies in the following sections.

### 2.3 Topology and inhibitory strength jointly shape degeneration responses

To quantify degeneration-induced changes in network activity, we evaluated complementary structural and dynamical observables. The mean firing rate (***λ***) captures overall activity level. Population synchrony was quantified using the Fano factor (***F F***), and spike-time irregularity using the coefficient of variation (***CV***) of inter-spike intervals.

We considered two synaptic weighting schemes (see Methods). In the *inhibition-boosted* weighting scheme shown in Fig. 2, excitatory synapses onto inhibitory neurons were stronger than excitatory synapses onto excitatory neurons, thereby preferentially boosting inhibitory recruitment. In the *standard* weighting scheme shown in Supplementary Fig. S2, excitatory synapses had the same strength regardless of whether their postsynaptic targets were excitatory or inhibitory, so the additional inhibitory bias present in the inhibition-boosted case was absent.

**Figure 2.**
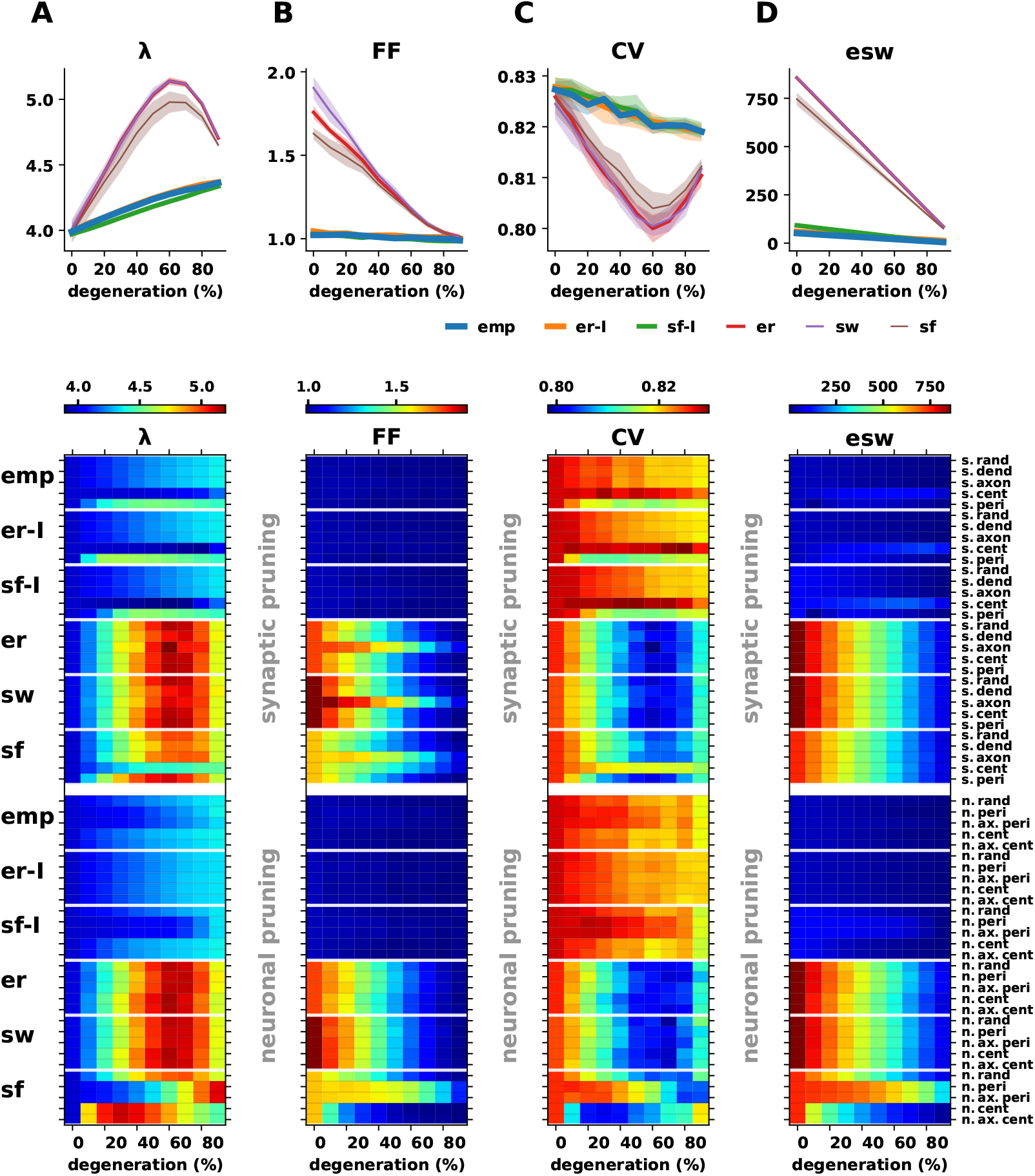
Degeneration responses depend jointly on network architecture and inhibitory strength. (A–D) Columns show mean firing rate (***λ***), population Fano factor (***F F***), ISI coefficient of variation (***CV***), and absolute effective synaptic weight (|***ESW*** |). Network classes are empirical (`emp`), Erdős–Rényi with inhibition-promoting architecture (`er-I)`, scale-free with inhibitory hubs (`sf-I`), Erdős–Rényi (`er`), small-world (`sw`), and scale-free (sf). **Top row:** Degeneration trajectories for synaptic pruning of random priority, corresponding to one slice of the heatmaps below. **Heatmaps:** Summary across pruning strategies for the tuned baseline condition (4.0 Hz), grouped by class, with synaptic pruning (upper panel) and neuronal pruning (lower panel). Under this moderate inhibition regime, networks with inhibition-promoting organization (`emp, er-I, sf-I`) exhibited relatively stable activity profiles over a wide range of degeneration, whereas networks without preferential organization of inhibitory connectivity (`er, sw, sf`) showed stronger deviations. This relative resilience was regime-dependent: increasing inhibitory strength reduced these differences and brought all classes toward similar dynamical responses (Supplementary Fig. S2).

We calibrated the intact networks to a common baseline firing rates to enable direct comparison of how degeneration altered activity dynamics across classes. Under the inhibition-boosted weighting scheme shown in Fig. 2, degeneration then induced clearly separated trajectories across network classes. This is most apparent in columns A–C: the empirical network and the two architectures with inhibition-promoting organization (`emp, er-I, sf-I`) remained close to their baseline operating point over a wide degeneration range, with only modest increases in firing rate (A), ***F F*** values staying near the asynchronous limit (B), and gradual changes in ***CV*** (C). In contrast, the remaining classes (`er, sw, sf`) showed much larger excursions in the same columns, including pronounced increases in firing rate and broader shifts in variability and irregularity. The key point was therefore not simply that some architectures were resilient and others were not, but that architectures differed in how much inhibitory scaling was required to keep degeneration trajectories close to the baseline regime.

This interpretation became especially clear when Fig. 2 was compared directly with Supplementary Fig. S2, which used the same color bars across classes and weighting schemes. Looking across the corresponding columns A–C in the two figures, the large deviations seen for `er, sw`, and `sf` under the inhibition-boosted comparison in Fig. 2 were strongly suppressed under the standard weighting scheme, and all six classes followed much more closely aligned trajectories. Thus, the latter three classes were not intrinsically non-resilient; rather, they required substantially larger inhibitory-to-excitatory synaptic strength ratios to achieve the same level of dynamical stability that `emp, er-I`, and `sf-I`already exhibited under the inhibition-boosted scheme. Put differently, inhibition-promoting architecture reduced the amount of inhibitory gain needed to tame degeneration-induced deviations.

The heatmaps in Fig. 2 further showed that this effect held across pruning strategies, not only for the random-pruning trajectories in the top row. Under the inhibition-boosted scheme, synaptic (Fig. 2 top-half) and neuronal pruning ((Fig. 2 bottom-half)) of `emp, er-I`, and `sf-I` generally remained confined to narrower color ranges, whereas `er, sw`, and `sf`explored much larger regions of the shared activity scale. Targeted perturbations still interacted with topology in meaningful ways. In hub-rich networks, especially `emp` and `sf-I`, central synaptic pruning (`s.cent`) tended to induce stronger changes than peripheral pruning (`s.peri`), consistent with the selective disruption of structurally important nodes and edges. Under neuronal pruning, differences between peripheral targeting based on total degree (`n.peri`) and out-degree (`n.ax.peri`) were also more pronounced in heterogeneous architectures than in homogeneous ones, indicating that degeneration responses remained sensitive to the detailed mode of structural removal even within the broader effect of inhibitory stabilization.

These comparisons showed that topology alone did not determine resilience. What mattered was how topology and inhibitory strength combined. Architectures in which inhibition was structurally reinforced were already stabilized under the inhibition-boosted weighting scheme, whereas architectures lacking that reinforcement could be brought back into the same resilient regime only when inhibition was made sufficiently strong under the standard scheme. The architectural advantage of `emp, er-I`, and `sf-I` therefore lay in requiring less inhibitory gain to preserve stable dynamics under degeneration, not in producing a qualitatively different end state once inhibition became strong enough.

To relate these dynamical changes to underlying structure, we introduced the Effective Synaptic Weight (ESW), defined as the average total synaptic input per neuron (node strength). ESW captured the combined effects of topology, synaptic strength, and excitation–inhibition balance. As expected, |**ESW**| decreased monotonically with degeneration in column D of Fig. 2 and Supplementary Fig. S2. Readers can see there that the ESW trajectories contract in a largely uniform way even when columns A–C still show appreciable differences in firing statistics. Similar ESW values could therefore still correspond to different levels of firing-rate stability and variability across classes, particularly under the inhibition-boosted scheme where architectural differences remained most visible. This indicated that while ESW provided a useful global summary of structural degradation, it did not by itself explain why some architectures required less inhibitory scaling than others to remain dynamically stable.

### 2.4 Weight-Aware Structural Features Reveal Structured Correlations with Network Dynamics

To examine how structural properties relate to network dynamics across diverse degeneration schemes and network classes, we computed pairwise Pearson correlations between structural descriptors and activity measures pooled across classes, pruning strategies, and degeneration stages (Fig. 3). We considered two categories of descriptors in this analysis: global and subpopulation-resolved. The global descriptors included mean degree ***k***, the fraction of shared presynaptic neighbours (***sh***), spectral radius (***specR***), and the absolute effective synaptic weight |**ESW**|, which summarize overall connectivity and coupling strength at the network level. The subpopulation-resolved descriptors captured mesoscale structure and included subpopulation-specific degrees (***k***_***XY***_), synaptic drive terms (***g***_***XY***_), their fluctuation scales (***σ***_***XY***_), and second-order interaction terms (***g***_***XYZ***_ **= *g***_***ZY***_ ***g***_***YX***_), where ***X, Y, Z* ∈ *{E, I}*** denote excitatory (***E***) and inhibitory (***I***) populations (e.g., ***g***_***EI***_ denotes coupling strength from inhibitory to excitatory subpopulations).

**Figure 3.**
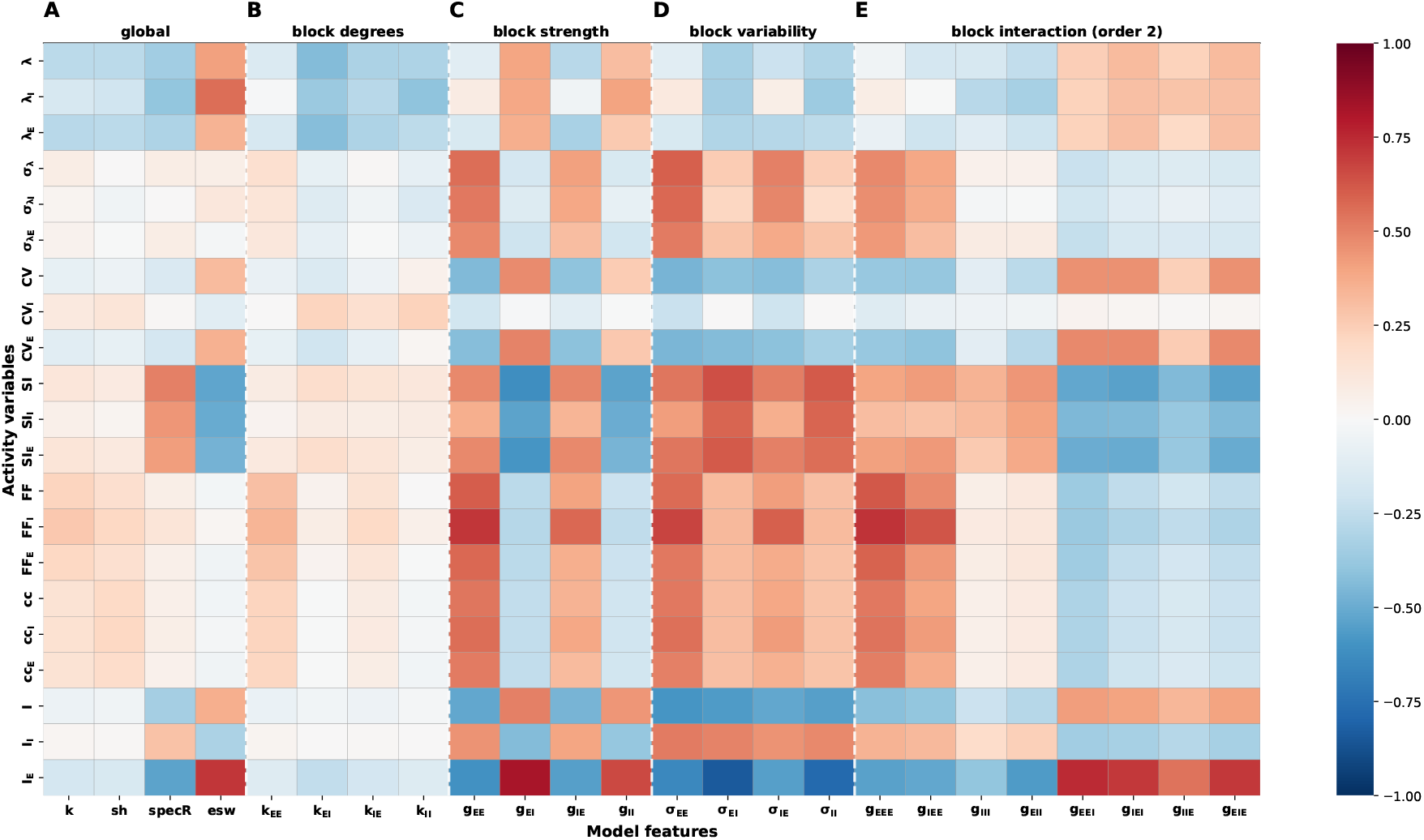
Structure–function correlations reveal dominant contributions of weight-aware subpopulation features. Heatmap of Pearson correlation coefficients between structural descriptors (columns) and activity measures (rows), pooled across network classes, pruning strategies, and degeneration stages under a fixed background-input configuration. Structural features are grouped as follows: **(A)** global descriptors including degree (***k***), shared presynaptic input (***sh***), spectral radius (***specR***), and effective synaptic weight (|**ESW**|); **(B)** subpopulation-resolved degrees ***k***_***XY***_ (***Y* →*X***), where ***X, Y* ∈ *{E, I}*** denote postsynaptic and presynaptic populations, respectively; **(C)** subpopulation synaptic drive terms ***g***_***XY***_ summarizing the mean synaptic coupling from population ***Y*** to ***X***; **(D)** subpopulation variability terms ***σ***_***XY***_ capturing fluctuations in subpopulation synaptic drive; and **(E)** second-order subpopulation interaction terms ***g***_***XYZ***_ corresponding to two-step population paths ***Z* →*Y* →*X***. Across activity variables, weight-aware subpopulation descriptors (***g***_***XY***_, ***σ***_***XY***_) and their interaction terms, ***g***_***XYZ***_, exhibit substantially stronger and more systematic correlations with firing rate, variability, and synchrony than purely topological quantities, indicating that mesoscale excitatory–inhibitory coupling structure is a primary determinant of network dynamics under degeneration.

The correlation matrix (Fig. 3) was organized with activity measures as rows and structural features as columns, grouped into global descriptors (A), subpopulation-resolved degrees (B), synaptic drive terms (C), fluctuation measures (D), and interaction terms (E). This layout enabled a direct comparison of how different classes of structural features related to distinct aspects of network activity.

A clear hierarchy emerged across these feature groups, with correlations becoming progressively stronger and more structured from purely topological descriptors to weight-aware coupling measures. The most important visual comparison in Fig. 3 is between panels A–B and panels C–E. In panels A–B, most entries remain close to zero or vary weakly across rows, indicating that connection counts alone carried limited information about degeneration-induced activity changes. Once synaptic weights were incorporated in panels C–E, broad and row-specific correlation patterns became much more pronounced.

Among the global descriptors in panel A, |**ESW**| and the spectral radius stood out from mean degree ***k*** and shared presynaptic input ***sh***. Their strongest signal appeared in the firing-rate and current-related rows, whereas ***k*** and ***sh*** remained weak almost everywhere. This was important because it showed that even at the global level, weight-aware summaries already outperformed purely topological ones. At the same time, the global descriptors were still too coarse to explain the full diversity of responses across subpopulations and observables.

Panel B showed that moving from global degrees to subpopulation-resolved degrees ***k***_***XY***_ improved interpretability but not yet explanatory power. The subpopulation-degree correlations were still comparatively weak and spatially diffuse, with only modest row-wise structure. Thus, subpopulation resolution by itself was not sufficient; the dominant gain came from incorporating synaptic weights.

The first major jump occurred in panel C, which displayed the subpopulation synaptic drive terms ***g***_***XY***_. Here the heatmap developed coherent vertical bands and much stronger row-wise structure. The clearest signal appeared for firing rates (***λ***) and synaptic currents (***I***), indicating that effective coupling at the subpopulation level captured the dominant control axis for average activity. Compared with panels A–B, panel C made clear that the network did not respond primarily to how many connections existed, but to how much weighted excitatory and inhibitory drive those connections delivered.

Panel D refined this picture further by showing that fluctuation features ***σ***_***XY***_ were particularly informative for variability-dominated observables. Their strongest structure appeared in the rows for firing-rate variability (***σ***_***λ***_), Fano factor (***F F***), spike-train irregularity (***CV***), and pairwise correlation (***cc***), whereas their relationship to mean firing rate (***λ***) was weaker than in panel C. This selective emphasis suggested that heterogeneity of synaptic input contributed more directly to dispersion, irregularity, and correlation structure than to the mean operating point itself.

The second-order interaction terms in panel E retained substantial structure but in a more motif-specific way. Rather than simply reproducing panel C, they showed differentiated patterns across synchrony, variability, and correlation-related rows, indicating that two-step pathways contributed information beyond mean subpopulation drive alone. This was especially useful for observables such as ***F F***, ***cc***, and several subpopulation-resolved measures, where the interaction panel remained visibly structured even when the degree panels did not.

Not all observables were captured equally well by the same feature family. Mean firing rates and synaptic currents aligned most clearly with the drive-based features in panels A and C, whereas variability and irregularity measures drew more strongly on panels D and E. Some rows, such as ***CV***_***I***_, showed weaker and more diffuse structure overall, indicating that certain subpopulation-specific quantities were less directly constrained by these coarse structural summaries.

Taken together, Fig. 3 showed that topology alone was insufficient to account for dynamical variation. What organized the heatmap was the progressive addition of weight information and subpopulation structure: global unweighted summaries were weak, subpopulation degrees alone remained limited, mean synaptic drives supplied the dominant signal, and fluctuation and interaction terms added more selective refinements. This motivated the use of weight-aware, subpopulation-resolved representations of network structure as a basis for subsequent analysis.

### 2.5 From global effective coupling to multivariate structural control of degeneration dynamics

Motivated by the correlation structure identified in Fig. 3, we next examined to what extent degeneration-induced dynamics can be explained by low-dimensional structural descriptors. We first considered the simplest weight-aware global quantity, the effective synaptic weight, **ESW =** _***X***,***Y***_ ***g***_***XY***_, as a univariate predictor of activity. By construction, **ESW** combines connection density, synaptic strength, and excitation–inhibition balance into a single scalar: its sign reflects whether the network is excitation- or inhibition-dominated, while its magnitude |**ESW**| quantifies deviation from balance.

#### Univariate structural control via effective synaptic weight

Figure 4 showed that degeneration trajectories were strongly organized by the absolute effective synaptic weight |**ESW**|, but that the quality of this organization depended on both the observable and the weighting scheme. In both panels, the clearest structure appeared for mean firing rate ***λ*** and net synaptic current ***I***_**syn**_: within each class, points aligned along relatively narrow, approximately linear trajectories as |**ESW**| changed. Thus, ESW captured a dominant axis of average dynamical change within classes.

**Figure 4.**
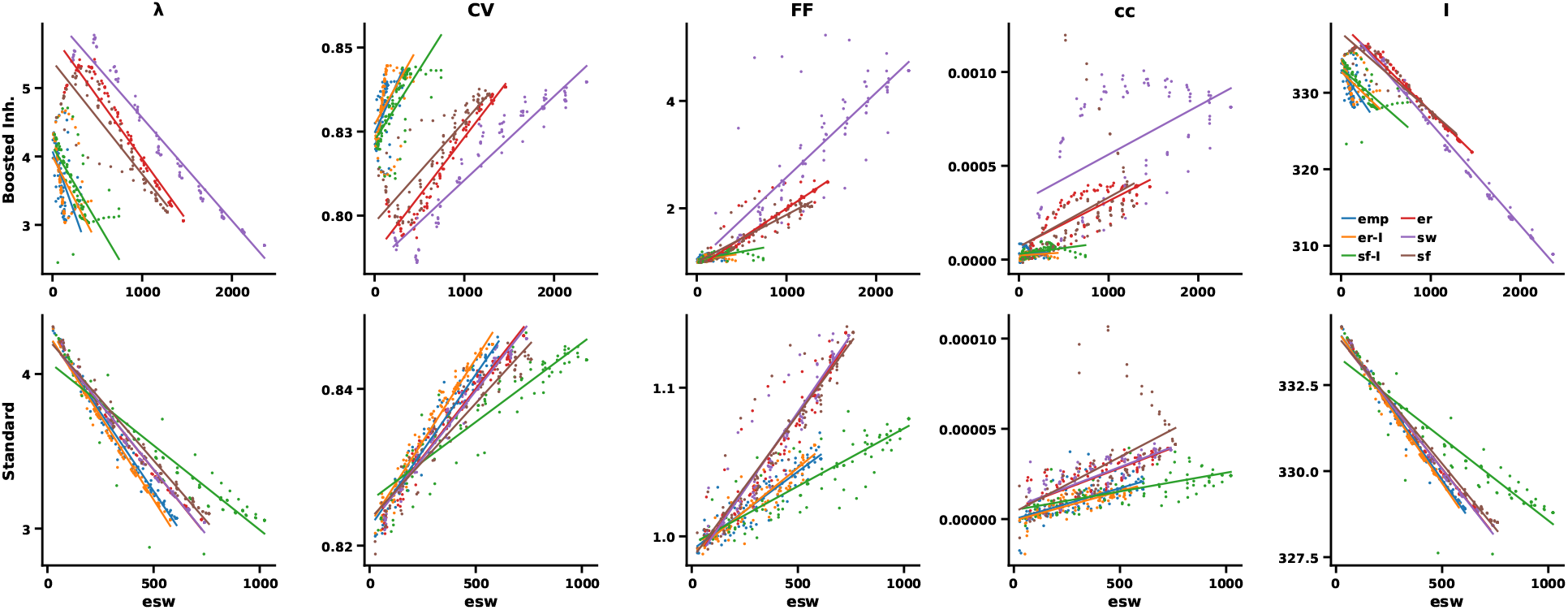
Effective synaptic weight organizes degeneration trajectories within classes but does not fully generalize across architectures. Scatter plots show mean firing rate (***λ***), spike-time irregularity (***CV***), population Fano factor (***F F***), and net synaptic current (***I***_**syn**_) as functions of absolute effective synaptic weight (|**ESW**|), pooled across degeneration stages and pruning strategies. Colors denote network classes (`emp, er-I, sf-I, er, sw, sf`); each point represents the mean over realizations for a given network state, and lines indicate class-specific degeneration trajectories in |**ESW**| space. **(A)** inhibition boosted weighting scheme. **(B)** Standard weighting scheme. Within each class, degeneration trajectories align along structured manifolds in |**ESW**| space, indicating strong coupling between total effective synaptic strength and activity measures. However, the mapping differs across network architectures, showing that |**ESW**| alone is insufficient as a universal predictor of activity across classes.

The limits of this univariate description were most visible in Fig. 4A. Under the inhibition-boosted weighting scheme, classes occupied clearly separated bands rather than collapsing onto a common curve. This was especially evident for ***λ***, where `sw` showed a much steeper dependence on |**ESW**| than `emp, er-I`, and `sf-I`, and for ***CV*** and ***F F***, where the class clusters remained visibly offset across much of the ESW range. In other words, similar changes in total effective coupling did not produce the same dynamical response in different architectures.

Figure 4B showed a partial collapse under the standard weighting scheme. The class trajectories became more tightly aligned, particularly for ***λ*** and ***F F***, indicating that stronger inhibition reduced some of the architecture-dependent spread. Even there, however, ***CV*** and ***cc*** retained appreciable dispersion at similar |**ESW**| values, showing that variability- and correlation-related observables still depended on structural detail beyond a single global coupling summary.

The comparison across panels therefore made the observable dependence clear: firing rates and synaptic currents were captured well by |**ESW**|, whereas irregularity, population variability, and pairwise correlation were not. ESW provided a useful univariate control axis, but the remaining class-dependent spread, particularly in ***CV***, ***F F***, and ***cc***, showed that a richer structural description was needed to account for architecture-specific deviations.

#### Multivariate structural control and predictive hierarchy

To capture topology-dependent deviations beyond the univariate description provided by |**ESW**|, we introduced a structured multivariate representation based on subpopulation-resolved coupling statistics. This representation includes subpopulation-level mean synaptic drives ***g***_***XY***_, fluctuation scales ***σ***_***XY***_, and second-order interaction terms ***g***_***XYZ***_ defined along directed population paths, yielding a 16-dimensional weight-aware feature space.

Using this representation, we predicted five activity observables (***λ, σ***_***λ***_, ***CV***, ***F F***, ***cc***) across all simulations using ridge regression with standardized features. Figure 5A–E showed that predicted and measured values lay close to the identity line for all five observables, with the tightest agreement for ***λ, σ***_***λ***_, and ***F F***, and broader but still structured scatter for ***CV*** and especially ***cc***. The summary curves in Fig. 5F,G further showed that this performance remained stable across test-set fractions and across a wide range of ridge penalties. In particular, the ***α* = 0** limit corresponding to ordinary least-squares regression performed almost identically to the mildly regularized ridge model, indicating that regularization was not the main source of predictive accuracy. Rather, the predictive signal was already present in the feature representation itself.

**Figure 5.**
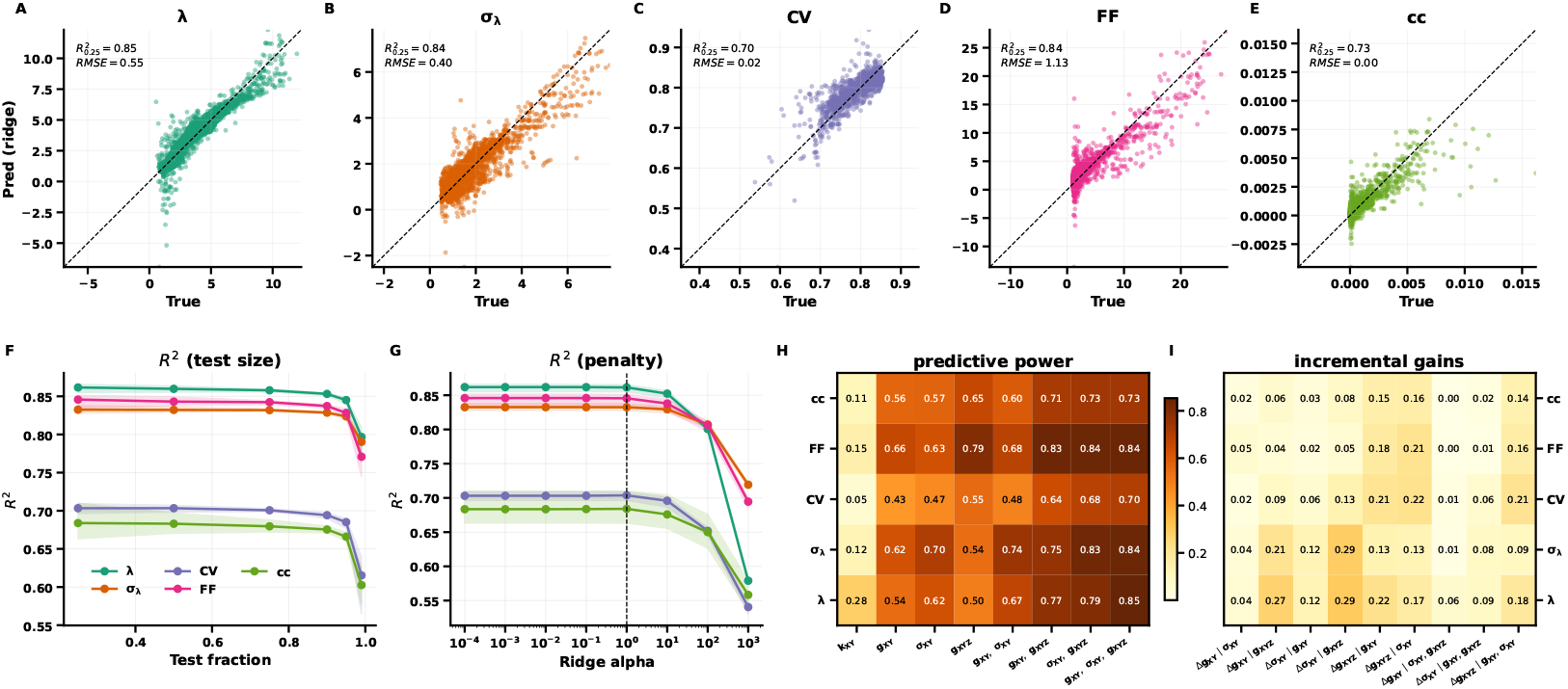
Multivariate structural features predict network activity and reveal a hierarchical contribution of feature groups. **(A–E)** Predicted versus measured values for mean firing rate (***λ***), firing-rate variability (***σ***_***λ***_), spike-time irregularity (***CV***), population Fano factor (***F F***), and pairwise spike-count correlation (***cc***), obtained using ridge regression on the same set of 16 weight-aware structural features. Predictions are shown for test data under repeated random train–test splits; dashed lines indicate identity. **(F)** Prediction accuracy (***R***^**2**^) as a function of test-set fraction, demonstrating stability across data splits. **(G)** Prediction accuracy as a function of ridge regularization strength, indicating robustness to the choice of penalty parameter. **(H)** Absolute predictive performance (***R***^**2**^) for models using different structural feature sets and their combinations, including subpopulation degrees (***k***_***XY***_), subpopulation synaptic drives (***g***_***XY***_), fluctuation scales (***σ***_***XY***_), and interaction terms (***g***_***XYZ***_). **(I)** Incremental gains (**Δ*R***^**2**^) when expanding feature sets. Here, **Δ*V***_**1**_ | ***V***_**2**_ **= *R***^**2**^**(*V***_**1**_ **∪ *V***_**2**_**) − *R***^**2**^**(*V***_**2**_) denotes the additional predictive power obtained by augmenting a model using feature set ***V***_**2**_ with features ***V***_**1**_. Large values indicate that ***V***_**1**_ contributes information not captured by ***V***_**2**_, whereas small values indicate substantial overlap between feature sets. Across all observables, predictions closely matched measured values and remained stable across test conditions. Weight-aware feature sets substantially outperformed degree-based models. Within that weight-aware family, subpopulation synaptic drives and fluctuation scales carried strongly overlapping predictive information, whereas interaction terms provided a smaller but more distinct additional contribution.

The heatmaps in Fig. 5H,I clarified where this predictive power came from. In panel H, the first column established the baseline limitation of subpopulation degrees alone: models based only on ***k***_***XY***_ performed worst across all observables. The remaining columns then compared progressively richer weight-aware descriptions, first as single feature families, then as pairwise combinations, and finally as the full three-family model. Panel H, in addition to separating weight-unaware and weight-aware models, showed how predictive power was distributed across the weight-aware feature families themselves. Subpopulation mean drives ***g***_***XY***_ and fluctuation features ***σ***_***XY***_ each supported strong prediction on their own, with ***σ***_***XY***_ often particularly informative for variability-related observables, while the best overall performance was typically reached when all three families were combined.

Panel I recast the same hierarchy in incremental form by asking what was gained when one feature family was added on top of another fit. This made clear that the contribution of a feature family depended on the context provided by those already included. In particular, adding ***g***_***XY***_ on top of ***σ***_***XY***_ or adding ***σ***_***XY***_ on top of ***g***_***XY***_ often yielded only modest gains, showing that these two families captured strongly overlapping aspects of the same underlying structure. By contrast, interaction terms ***g***_***XYZ***_ more often retained a distinct incremental effect once the lower-order weight-aware descriptors were already in the model. This interpretation was reinforced by the coefficient analysis in Supplementary Fig. S3. Across the orthogonalized decompositions, ***g***_***XY***_ and ***σ***_***XY***_ encoded largely overlapping predictive information: whichever one was entered first captured much of the variance available to the other. Interaction terms, by contrast, typically retained a clearer incremental contribution after the lower-order features had already been accounted for, indicating that they captured pathway-dependent structure not fully absorbed by the mean and fluctuation summaries.

At the same time, the multivariate description was not strongly compressible to just a few latent axes. A principal-component analysis of the standardized 16-dimensional feature matrix showed an effective rank of 15, and predictive performance for all five observables reached its maximum at 15 principal components while already lying within 0.01 of that maximum by 14 components. Consistent with this, the first 14 principal components explained 99.998% of the total variance, whereas the 15th component restored the remaining numerically non-redundant direction. The multivariate model should therefore be viewed as compact but effectively near-full-dimensional, rather than low-dimensional reduction.

The comparison between the univariate global |**ESW**| model and the multivariate subpopulation-resolved model clarified both the advantages of the richer representation and the limits of its generalization. As shown in Supplementary Fig. S4, the multivariate representation consistently outperformed |**ESW**| within classes, demonstrating that the added descriptors ***g***_***XY***_, ***σ***_***XY***_, and ***g***_***XYZ***_ captured substantial structure that the univariate summary missed. At the same time, leave-one-network-class-out transfer remained class-dependent (Supplementary Fig. S4D): transfer was strong for `er, sw`, and `sf`, but degraded markedly for `sf-I`. Thus, the richer structural representation captured much more variance than ESW alone without becoming fully architecture-agnostic.

Overall, degeneration dynamics were described most effectively by a compact, multivariate subpopulation-resolved representation that extended the univariate ESW picture. The main message was not that one particular lower-order feature family dominated all others, nor that ridge regularization created the predictive gain. Rather, resolving synaptic coupling at the subpopulation level was essential, mean drives and fluctuation scales carried largely overlapping predictive content, interaction motifs supplied an additional refinement, and the resulting representation captured architecture-dependent variability far better than |**ESW**| alone.

### 2.6 Degeneration Schemes Trace Structured Trajectories in Activity Manifold Space

So far, we had analyzed the evolution of structural and activity features along degeneration individually. However, degeneration affected all these features jointly, in ways that depended on network class and degeneration strategy. To obtain a global view of how degeneration reshaped network activity beyond individual metrics—and to reveal latent directions of variation that might discriminate among distinct degeneration schemes—we embedded network states into a low-dimensional activity manifold using t-distributed stochastic neighbor embedding (t-SNE) applied to the full multivariate activity profile. Each point represented a network state at a given degeneration stage, and proximity reflected similarity across all activity measures jointly.

Figure 6 provided a geometric summary of the multivariate activity space introduced in the previous sections. In panels A–D, the same t-SNE embedding was recolored by stage, network class, pruning strategy, and weighting scheme. The most robust pattern was the stage gradient in panel A: within many local branches of the embedding, early stages and late stages were arranged smoothly rather than being intermixed, indicating that degeneration moved network states along structured directions in activity space rather than scattering them randomly.

**Figure 6.**
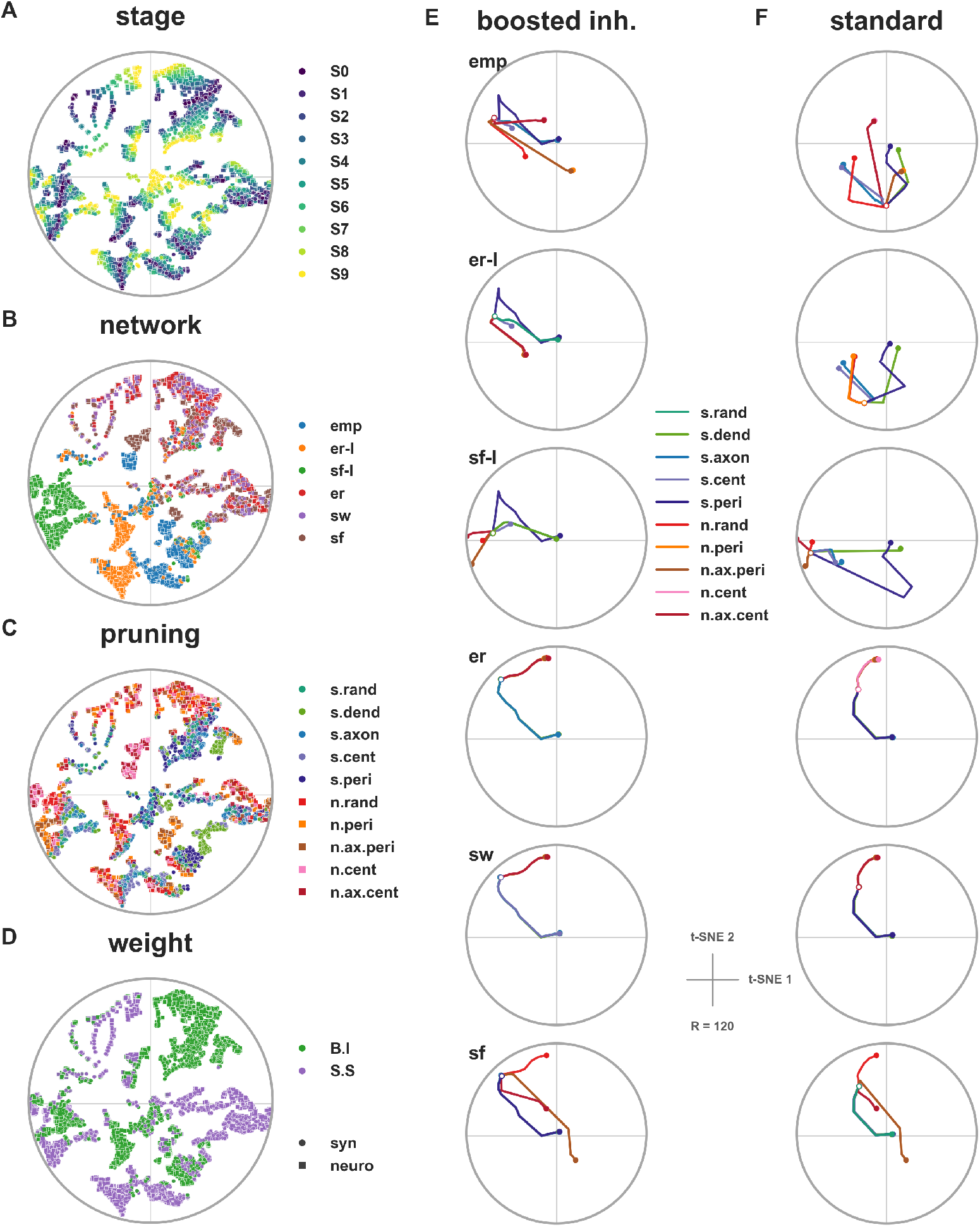
Degeneration trajectories in multivariate activity space. Two-dimensional t-SNE embedding of network states constructed from their full multivariate activity profiles. **(A–D)** The same embedding colored by different experimental factors: degeneration stage, network class, pruning strategy, and synaptic weighting scheme, respectively. The embedding organizes network states into coherent regions of activity space rather than random dispersion. **(E–F)** Degeneration trajectories within each network class, with successive stages connected by lines. Column (E) shows the inhibition-boosted weighting scheme and column (F) the standard weighting scheme. Across classes, trajectories evolve smoothly with degeneration stage, while their geometry depends on both architecture and weighting scheme. Network identity and synaptic weighting create the clearest large-scale structure in the embedding, whereas pruning strategy acts as a secondary source of variation that modulates trajectory shape within each class.

Panels B and D showed that network identity and weighting scheme were the dominant large-scale organizers of this embedding. In panel B, classes occupied partially separated regions, with substantial overlap among the inhibition-promoting architectures (`emp, er-I, sf-I`) and clearer separation of `er, sw`, and `sf` into other parts of the map. Panel D showed an additional split by weighting scheme, indicating that the choice between inhibition-boosted and standard synaptic weighting reshaped the overall geometry of the activity manifold.

By contrast, panel C suggested that pruning strategy acted mainly as a secondary source of structure. Different pruning rules did modulate where points fell within the embedding, but they did not form a clean global partition comparable to that seen for stage, class, or weighting.

The trajectory panels in Fig. 6 E,F made these comparisons more concrete by connecting successive stages within each class. Across all six classes, degeneration followed smooth, low-dimensional paths rather than erratic jumps, but the geometry of those paths differed markedly between the two weighting schemes. In the inhibition-boosted case (E), trajectories spanned a broader portion of the manifold and showed clearer class-to-class differences. In the standard case (F), trajectories became more compressed and more similarly oriented, consistent with the stronger cross-class alignment already seen in Supplementary Fig. S2 and Fig. 4B.

The comparison across classes also revealed a clear grouping. In panel E, the empirically organized and inhibition-promoting architectures (`emp, er-I, sf-I`) followed relatively compact trajectory fans that remained closer to one another in both location and orientation, whereas `er, sw`, and `sf` showed larger excursions and stronger divergence across pruning rules. This mirrored the earlier observation that the first group was already stabilized under the inhibition-boosted weighting scheme, while the latter group required stronger effective inhibition to achieve comparable alignment.

Within this latter group, `er` and `sw` were especially similar. In both panels E and F, their trajectories occupied neighboring parts of the embedding and followed closely related directions of drift, consistent with their comparatively homogeneous degree structure and lack of inhibitory hub enrichment. The scale-free network (`sf`) shared some aspects of this broader group but was less tightly matched to `er` and `sw`, likely because its global hub heterogeneity still introduced additional curvature and spread.

The clearest qualitative contrast therefore lay not in a universal separation of synaptic versus neuronal pruning, but in how different architectures traversed the same activity space under different inhibitory regimes. In some classes, synaptic and neuronal pruning families occupied somewhat different parts of the local trajectory fan, whereas in others they remained close. The embedding thus supported a cautious interpretation: degeneration mode and pruning strategy affected trajectory shape, but the strongest structure reflected architecture and weighting scheme.

Taken as a whole, Fig. 6 reinforced the view that degeneration responses were constrained to a shared low-dimensional activity manifold shaped primarily by inhibitory gain and network architecture. The manifold representation did not replace the scalar structural analyses; rather, it complemented them by showing how the same underlying control variables organized the geometry of multivariate activity change across classes and degeneration conditions.

These findings showed that vulnerability and resilience to structural damage emerged from the interplay between inhibitory gain, subpopulation organization, and recurrent coupling geometry.

## Discussion

The present results point to a view of degeneration in which structural loss does not translate directly into dynamical failure. Instead, damage is filtered through the circuit mechanisms that normally stabilize recurrent activity. The same amount of synaptic or neuronal removal can therefore produce mild drift, strong rate elevation, or altered variability depending on how inhibitory control is organized. This is the main sense in which resilience should be read here: not as resistance to anatomical loss, but as the ability of the remaining circuit to keep activity near a health-like asynchronous-irregular operating point while its structure is progressively degraded.

This perspective gives the abstract pruning strategies a biological role. Neurodegenerative processes rarely remove connections as a uniform random thinning; structural loss is shaped by molecular pathology, synaptic state, cell-type vulnerability, and the circuit context in which damage occurs. Alzheimer’s disease illustrates this point. Synaptic dysfunction is an early and prominent component of the disease, and both amyloid-***β*** and tau have been linked to synapse loss. Experimental work further suggests that immune-mediated mechanisms, including complement and microglial activity, can contribute to early synapse elimination in disease models [23, 24]. Although tau is primarily intracellular, the regional progression of tau pathology appears to follow connected neural systems, suggesting that synaptic dysfunction and circuit architecture may interact during disease progression [25, 26]. Similar selectivity is not limited to Alzheimer’s disease: projection neurons, striatal medium spiny neurons, and inhibitory interneuron populations are affected differently across disorders and disease stages [27, 28]. The pruning rules used here should therefore not be interpreted as literal models of specific neurodegenerative diseases. They are controlled abstractions of the broader fact that degeneration is biased by biological and structural context rather than being a purely random loss process.

The strongest implication concerns inhibition. A large body of work shows that inhibition is not merely a counterweight to excitation, but a stabilizing feedback system that shapes gain, variability, correlations, and the transition between asynchronous and coherent regimes. Inhibition-stabilized and stabilized supralinear network theories show that recurrent excitation can be held in a controlled regime only when feedback inhibition is sufficiently strong and properly recruited [29, 30, 31]. Our results extend this idea to progressive structural loss. A circuit can have the same nominal E/I composition, or even the same total effective coupling, and still respond differently to degeneration if inhibitory neurons occupy different structural positions. Thus, E/I balance by itself is an incomplete descriptor for a degenerating network: what matters is not only how much inhibition is present, but whether the architecture allows inhibition to be engaged at the points where recurrent drift is generated.

This also clarifies the relation between architecture and inhibitory strength. The empirical, `er-I`, and `sf-I` networks were not resilient because they belonged to an entirely separate dynamical category. Rather, their organization made inhibition more efficient. Canonical random, small-world, and scale-free networks could also be stabilized when inhibitory strength was increased, but they required more gain to reach the same qualitative regime. Architecture therefore acts like a gain-efficiency factor: inhibitory hubs, enriched excitatory drive onto inhibitory populations, or preserved E/I subpopulation densities reduce the synaptic strength needed to keep activity bounded during degeneration. This distinction is important biologically because it separates two routes to resilience: increasing inhibition globally, or preserving the structural pathways through which inhibition is recruited.

This inhibition-centered interpretation motivated the structural prediction analysis. Because all baseline networks were calibrated in an inhibition-dominated regime, global effective synaptic weight (ESW) was negative and served as a simple proxy for effective inhibitory dominance. It was therefore a natural first predictor of degeneration-induced activity changes: if resilience reflected inhibitory control, then a global weighted-coupling summary should capture part of the dynamical drift. This univariate ESW model was informative because it organized firing rate and synaptic current well within network classes, showing that degeneration changed activity along a shared coupling axis. However, ESW did not collapse all architectures onto a single curve: similar global ESW values could still support different variability, irregularity, and synchrony. The multivariate model kept the same principle –weighted connectivity as structural control– but resolved it at a finer scale. Instead of a single global sum, it used subpopulation-resolved weights, fluctuations in those weights, and second-order interaction terms, making explicit which weighted E/I interactions remain invisible to a single global measure. This finer model substantially mitigated the limitations of the ESW-only view and improved prediction across activity measures.

The low-dimensional projection of multivariate activity space should be interpreted as a descriptive summary rather than a mechanistic state space. Within that view, network architecture, degeneration stage, and inhibitory regime provided the clearest large-scale organization, whereas pruning rule and synaptic versus neuronal degeneration mainly shaped local trajectory patterns. This suggests that different microscopic degeneration processes may converge onto similar coarse activity changes, while still leaving more subtle differences in the joint profile of rate, variability, and synchrony. Such convergence is consistent with the idea that EEG, MEG, or fMRI markers can reveal broad network dysfunction while remaining limited in their ability to identify the underlying cellular process without additional structural or longitudinal constraints [32, 33].

One limitation of the present model is that it treats degeneration as removal without repair. Biological circuits respond to damage through homeostatic plasticity, synaptic scaling, dendritic remodeling, and sprouting. Such mechanisms can be stabilizing, but the present results suggest why compensation may also fail. Adding or strengthening connections is not equivalent to restoring the original architecture. If compensatory sprouting creates connections semi-randomly, or if homeostatic scaling boosts activity without preserving inhibitory routing, it may recover total input while blurring the fine organization that made inhibition efficient. In that case, the circuit could look repaired by coarse connectivity measures while still drifting in variability, synchrony, or susceptibility to later perturbation.

Several further simplifications should be kept in mind. The simulations used leaky integrate-and-fire neurons and were calibrated to weakly active irregular regimes, so they do not address transitions into epileptiform, oscillatory, or strongly synchronized pathological states. The architectures were stylized comparisons around one empirical microcircuit rather than disease-specific reconstructions. The pruning rules were intentionally abstract, and the low-dimensional activity projection is a descriptive summary of model trajectories rather than a direct biological state space. These choices made it possible to isolate architecture, inhibitory gain, and degeneration mode, but future work should test whether the same control principles hold under cell-type-specific physiology, plasticity, spatial constraints, and disease-specific molecular mechanisms.

Overall, the study suggests that circuit resilience under degeneration is a graded property of inhibitory organization. Degeneration can be diverse at the microscopic level, yet still produce partially shared macroscopic trajectories because all damage is filtered through the same recurrent stabilizing machinery. Conversely, similar gross levels of damage can lead to distinct activity outcomes when inhibitory recruitment is organized differently. This provides a bridge between structural degeneration, population-level biomarkers, and mechanistic prediction: the key question is not only what is lost, but whether the remaining circuit can still recruit inhibition through the pathways that keep recurrent activity under control.

## 3 Methods

### Networks

We studied six network classes spanning one empirical reference class and five synthetic comparison classes. Throughout, we used the adjacency convention ***A*[tgt, src] = 1** (columns are sources, rows are targets), and all models produced directed adjacency matrices with zero diagonal.

#### Empirical connectivity (emp)

Structural connectivity was derived from the anatomically constrained statistical connectome of Landau et al. [15], in which pairwise connection likelihoods were estimated from three-dimensional soma distributions and axo-dendritic overlap statistics in layer 4 barrel cortex. The resulting contact matrix ***A***_***ij***_ encodes the expected number of anatomical contacts between presynaptic neuron ***j*** and postsynaptic neuron ***i***, thereby capturing structured heterogeneity and excitatory–inhibitory in-degree correlations.

To obtain binary networks with controlled sparsity, we generated adjacency matrices using a shifted Bernoulli sampling scheme:

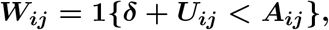

where ***U***_***ij***_ **~ *U* (0, 1)** and ***δ*** is a density-control parameter. This is equivalent to sampling connections with probability ***P***_***ij***_ **= max(0, *A***_***ij***_ **− *δ*)**, effectively suppressing weak anatomical contacts while preserving the relative heterogeneity of stronger projections. The resulting empirical networks contained 3611 neurons in total, comprising 2931 excitatory and 680 inhibitory neurons (an E:I ratio of approximately 4.3:1). We set ***δ* = 0.15**, yielding networks with approximately **10%** connection density without self-connections.

This empirical class served as the reference scaffold for generating synthetic comparison classes that preserved selected global constraints, in particular the total synapse count, while varying degree heterogeneity and mesoscale excitatory–inhibitory organization.

#### Erdős–Rényi (er)

This class was generated as a homogeneous Erdős–Rényi graph with fixed total edge count, providing a maximally random reference with narrow degree variability and no imposed mesoscale structure.

#### Small-world (sw)

This class was generated as a small-world graph obtained by applying limited rewiring to a regular ring lattice, together with interleaved inhibitory relabeling, thereby preserving substantial local regularity while introducing short-cuts.

#### Scale-free (sf)

This class was generated using a scale-free construction approximating a powerlaw degree distribution with exponent between 2 and 3, with hubs distributed across both excitatory and inhibitory populations.

#### Erdős–Rényi with boosted inhibition architecture (er-I)

This class was generated as a subpopulation-constrained Erdős–Rényi construction that preserves the empirical subpopulation connection densities (equivalently, edge counts) within and between populations, i.e., the four subpopulations ***E* → *E, E* → *I, I* → *E***, and ***I* → *I***, while randomizing connections within each subpopulation pair (see Table 1).

**Table 1.** Subpopulation-level connection probabilities **[*p***_***EE***_, ***p***_***EI***_, ***p***_***IE***_, ***p***_***II***_] for each network class.

| Ensemble | $p_{II}$ | $p_{EI}$ | $p_{IE}$ | $p_{EE}$ |
| --- | --- | --- | --- | --- |
| emp | 0.149 | 0.179 | 0.117 | 0.087 |
| er-I | 0.149 | 0.179 | 0.117 | 0.087 |
| sf-I | 0.327 | 0.162 | 0.161 | 0.071 |
| er | 0.108 | 0.108 | 0.108 | 0.108 |
| sw | 0.108 | 0.108 | 0.108 | 0.108 |
| sf | 0.106 | 0.107 | 0.107 | 0.108 |

#### Scale-free with boosted inhibition architecture (sf-I)

This class used the same underlying scale-free construction as sf, but differed in node labeling: inhibitory neurons were given priority to occupy hub positions, thereby enriching the inhibitory population in high-degree nodes.

### Degeneration schemes and pruning strategies

Degeneration was applied at two levels: synaptic (edge-level) and neuronal (node-level). In both cases, degeneration progressed sequentially in nine stages beyond the intact network, with each stage removing **10%** of the remaining elements from the previous stage. Pruning was performed in a balanced manner across excitatory and inhibitory populations.

#### Sequential pruning as masking

Let ***A***^**(0)**^ **∈ *{*0, 1*}***^***N×N***^ denote the intact adjacency (zero diagonal). For synaptic pruning we define an edge-survival mask 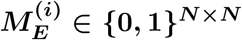 and set

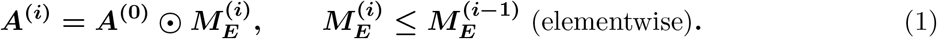

For weighted connectivity ***W*** ^**(0)**^, the pruned weighted matrix is 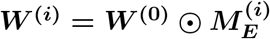.

For neuronal pruning we define a node-survival vector 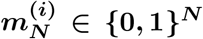 and its diagonal matrix 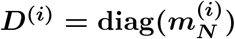, yielding

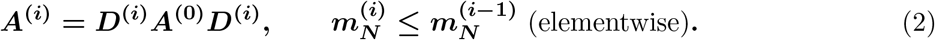

This operation removes all in- and out-edges of deleted neurons simultaneously.

#### Synaptic degeneration (edge-level)

Synaptic degeneration was implemented as sequential edge removal applied to the adjacency matrix (Eq. 1). At each stage, 10% of the remaining synapses were removed in a population-balanced manner across excitatory and inhibitory subpopulations.

The strategies differed in how candidate edges were ranked for pruning priority. Two complementary ranking principles were used: degree-based ranking and maximum-matching–based ranking.

#### Degree-based ranking

These strategies prioritized edges according to local connectivity statistics of the associated neurons. Specifically, edges could be ranked based on the in-degree of the target neuron (dendritic perspective), the out-degree of the source neuron (axonal perspective), or without regard to connectivity structure (random selection).

#### Maximum-matching–based ranking

The remaining strategies were based on a structural decomposition derived from successive maximum matchings of the directed network. Maximum matchings play a central role in structural controllability theory, where they determine the minimum set of driver nodes required to control a network [34, 35]. To obtain an ordering of edges, we computed successive maximum matchings on a bipartite representation of the directed graph using the Hopcroft–Karp algorithm. At each iteration a maximum matching ***M***_***i***_ was extracted from the remaining edge set, yielding a decomposition 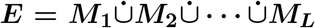. Because a matching contains no edges sharing the same source or target node, each node could contribute at most one outgoing and one incoming edge per layer. Consequently, edges incident to low-degree nodes tended to be exhausted in early layers, whereas edges associated with high-degree nodes persisted across many matching layers. This property induced a structural ranking of edges according to their persistence under the matching constraint. Pruning could therefore proceed either from early layers, removing edges associated with nodes that became isolated quickly, or from late layers, trimming edges that remained supported by highly connected nodes.

Using these ranking principles, we defined five pruning strategies:

- **Random pruning (**`s.rand`). Edges were removed uniformly at random.
- **Dendritic pruning (**`s.dend`). Edges were prioritized according to the in-degree of their target neuron, emphasizing pruning of highly connected dendritic targets.
- **Axonal pruning (**`s.axon`). Edges were prioritized according to the out-degree of their source neuron, emphasizing pruning of strong broadcasting neurons.
- **Peripheral pruning (**`s.peri`). Edges were removed in the order ***M***_**1**_, ***M***_**2**_, …, prioritizing edges associated with nodes that exhausted their connectivity early in the matching decomposition.
- **Central pruning (**`s.cent`). Edges were removed in the reverse order ***M***_***L***_, ***M***_***L*−1**_, …, preferentially trimming edges associated with nodes that remained connected across many matching layers (typically hubs).

#### Neuronal degeneration (node-level)

Neuronal degeneration was implemented as sequential node removal using the masking operation in Eq. 2. We considered five pruning strategies: random neuronal pruning, peripheral pruning, axonally peripheral pruning, central pruning, and axonally central pruning. At each stage, 10% of the remaining neurons were removed, balanced across excitatory and inhibitory populations.

The strategies differed in how neurons were ranked prior to removal:

- **Random neuronal pruning (**`n.rand`). randomly sampled neurons were targeted.
- **Peripheral pruning (**`n.peri`). weakly connected neurons were prioritized.
- **Axonally peripheral pruning (**`n.ax.peri`). weak broadcasters (low out-degree) were prioritized.
- **Central pruning (**`n.cent`). hub neurons were prioritized.
- **Axonally central pruning (**`n.ax.cent`). strong broadcasters (high out-degree) were prioritized.

### Weighting schemes

Synaptic weights were assigned according to the presynaptic and postsynaptic population identities while preserving global network connection density. For each directed synapse from a presynaptic population ***Y* ∈ *{E, I}*** to a postsynaptic population ***X* ∈ *{E, I}***, we denote its synaptic weight by ***w***_***XY***_ (i.e., ***Y* → *X***). Two weighting schemes were considered: *boosted-inhibition scheme* and *standard setting scheme*. In the boosted-inhibition scheme, excitatory synapses onto inhibitory neurons were stronger than excitatory synapses onto excitatory neurons, i.e., ***w***_***IE***_ ***> w***_***EE***_, thereby preferentially recruiting inhibition. In the standard scheme, excitatory synapses had the same strength regardless of whether their postsynaptic targets were excitatory or inhibitory, i.e., ***w***_***IE***_ **= *w***_***EE***_

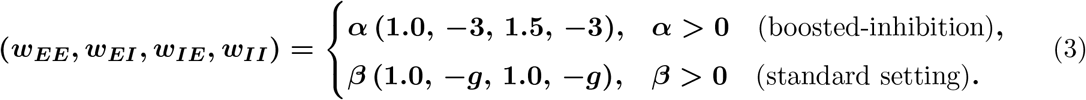

In the boosted-inhibition scheme, ***α*** scales all synaptic weights while maintaining the relative enhancement of excitatory synapses onto inhibitory targets. In the standard scheme, ***β*** sets the common excitatory scale and ***g*** sets the inhibitory-to-excitatory weight ratio.

### Experimental design and datasets

We analyzed two dataset families, which we refer to as *untuned* and *tuned*. Each family contains two subsets corresponding to the two weighting schemes introduced above: the boosted-inhibition scheme and the standard scheme. We use the term *untuned* for datasets in which synaptic scaling parameters were swept without enforcing a common baseline activity level across network classes, and *tuned* for datasets in which synaptic scaling parameters were selected to match prescribed baseline firing-rate targets across classes.

Across both dataset families, we varied synaptic-weight scaling parameters while keeping the background input configuration fixed. Both dataset families were generated using the same background input configuration, corresponding to a Poisson drive of 8 kHz with connection weight 7.69.

For the untuned datasets, the inhibitory ratio was fixed to ***g* = 5** in the standard subset. In each subset, the corresponding synaptic scaling parameter was sampled at ten equally spaced values between 0.1 and 15.

For the tuned datasets, synaptic scaling factors were selected to produce comparable baseline population firing rates across network classes. In the standard subset, a stronger inhibitory ratio had to be used, particularly to place the `er, sw`, and `sf` classes in a shared target firing-rate regime with the real connectivity data. We therefore chose ***g* = 10**, twice the untuned value, for all network classes in the standard subset for consistency, even though the empirical and subpopulation-preserving Erdős–Rényi classes (`emp, er-I`) did not require this stronger ratio. Ten target rates were chosen equally spaced between 2.5 and 4.2 Hz. For stage-0 networks, we first evaluated twenty equally spaced values of the synaptic scaling parameter in each weighting-scheme subset, using the interval 0.3 to 15 for the boosted-inhibition scheme and 0.03 to 3 for the standard scheme. Exponential fits were then used to approximate the relationship between scaling parameter and baseline firing rate for each network class, allowing us to estimate the preimage of the ten target rates. Table 3 lists the resulting fitted scaling parameters for each network class and target firing rate.

**Table 2.**
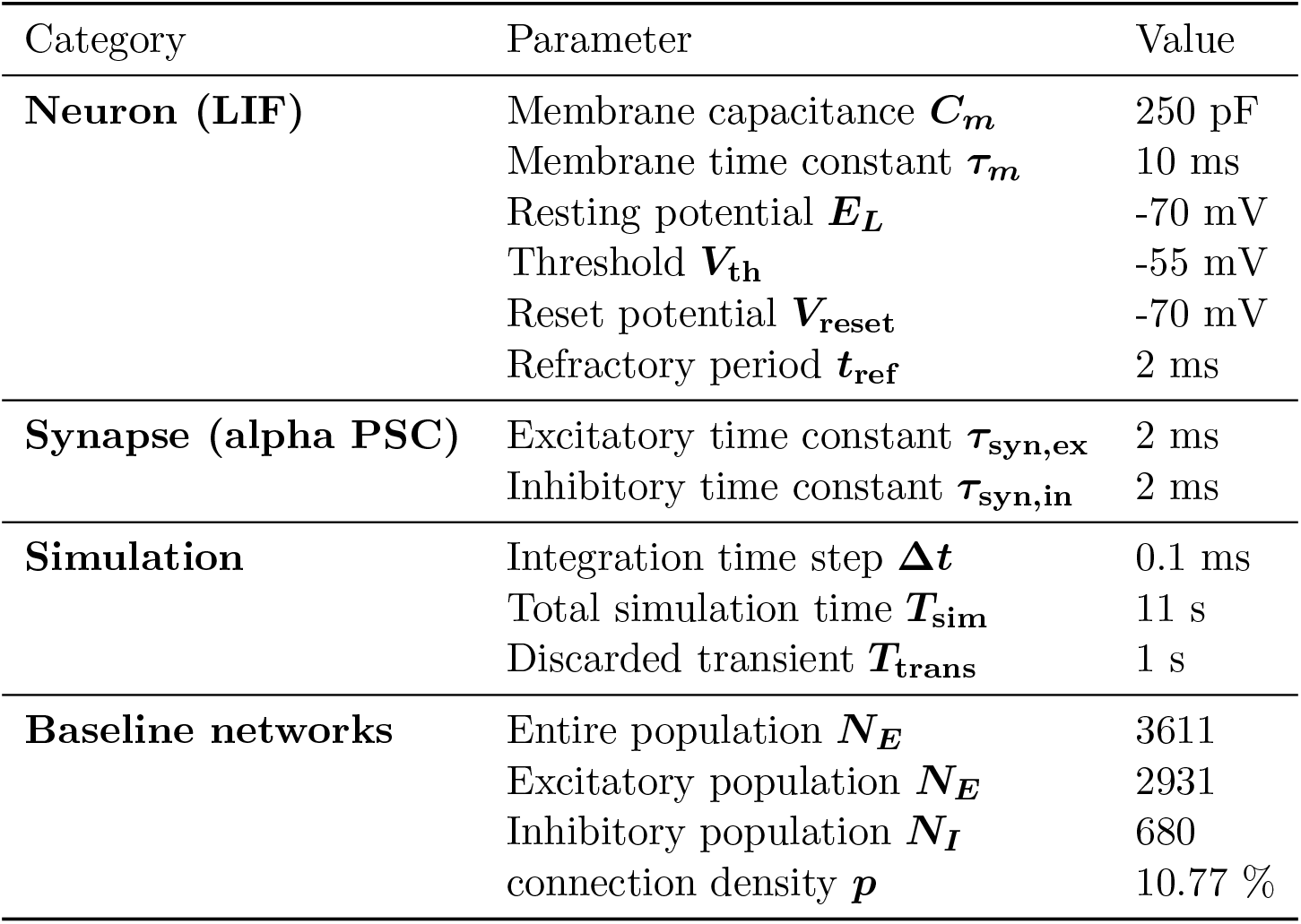
Neuron, synapse, and simulation parameters.

**Table 3.** Fitted scaling parameters for the tuned dataset. Rows correspond to target baseline firing rates (Hz). Columns under ***α*** give the fitted boosted-inhibition scaling parameter for each network class, and columns under ***β*** give the fitted standard-scheme scaling parameter for each network class.

| Target rate | $\alpha$ (boosted-inhibition) | | | | | | $\beta$ (standard) | | | | | |
| --- | --- | --- | --- | --- | --- | --- | --- | --- | --- | --- | --- | --- |
|  | emp | er-I | sf-I | er | sw | sf | emp | er-I | sf-I | er | sw | sf |
| 4.20 | 2.361 | 2.101 | 8.362 | 15.000 | 4.079 | 8.230 | 0.633 | 0.570 | 1.928 | 1.981 | 1.385 | 2.284 |
| 4.01 | 2.034 | 1.852 | 7.367 | 15.000 | 3.297 | 6.694 | 0.518 | 0.474 | 1.383 | 1.494 | 0.966 | 1.699 |
| 3.82 | 1.743 | 1.619 | 6.516 | 13.857 | 2.704 | 5.712 | 0.415 | 0.389 | 1.129 | 1.132 | 0.690 | 1.215 |
| 3.63 | 1.480 | 1.420 | 5.805 | 9.405 | 2.203 | 5.015 | 0.336 | 0.320 | 0.871 | 0.886 | 0.485 | 0.926 |
| 3.44 | 1.260 | 1.223 | 5.065 | 7.131 | 1.748 | 4.290 | 0.274 | 0.261 | 0.669 | 0.676 | 0.350 | 0.681 |
| 3.26 | 1.088 | 1.045 | 4.665 | 6.109 | 1.359 | 3.733 | 0.209 | 0.207 | 0.505 | 0.509 | 0.250 | 0.498 |
| 3.07 | 0.886 | 0.854 | 4.160 | 4.954 | 1.065 | 3.329 | 0.161 | 0.158 | 0.375 | 0.383 | 0.180 | 0.367 |
| 2.88 | 0.710 | 0.692 | 3.797 | 4.414 | 0.768 | 2.923 | 0.116 | 0.111 | 0.255 | 0.263 | 0.118 | 0.251 |
| 2.69 | 0.524 | 0.532 | 3.409 | 3.867 | 0.546 | 2.489 | 0.072 | 0.075 | 0.169 | 0.169 | 0.072 | 0.154 |
| 2.50 | 0.328 | 0.327 | 3.057 | 3.417 | 0.305 | 2.150 | 0.040 | 0.037 | 0.083 | 0.083 | 0.034 | 0.074 |

**Table 4.**
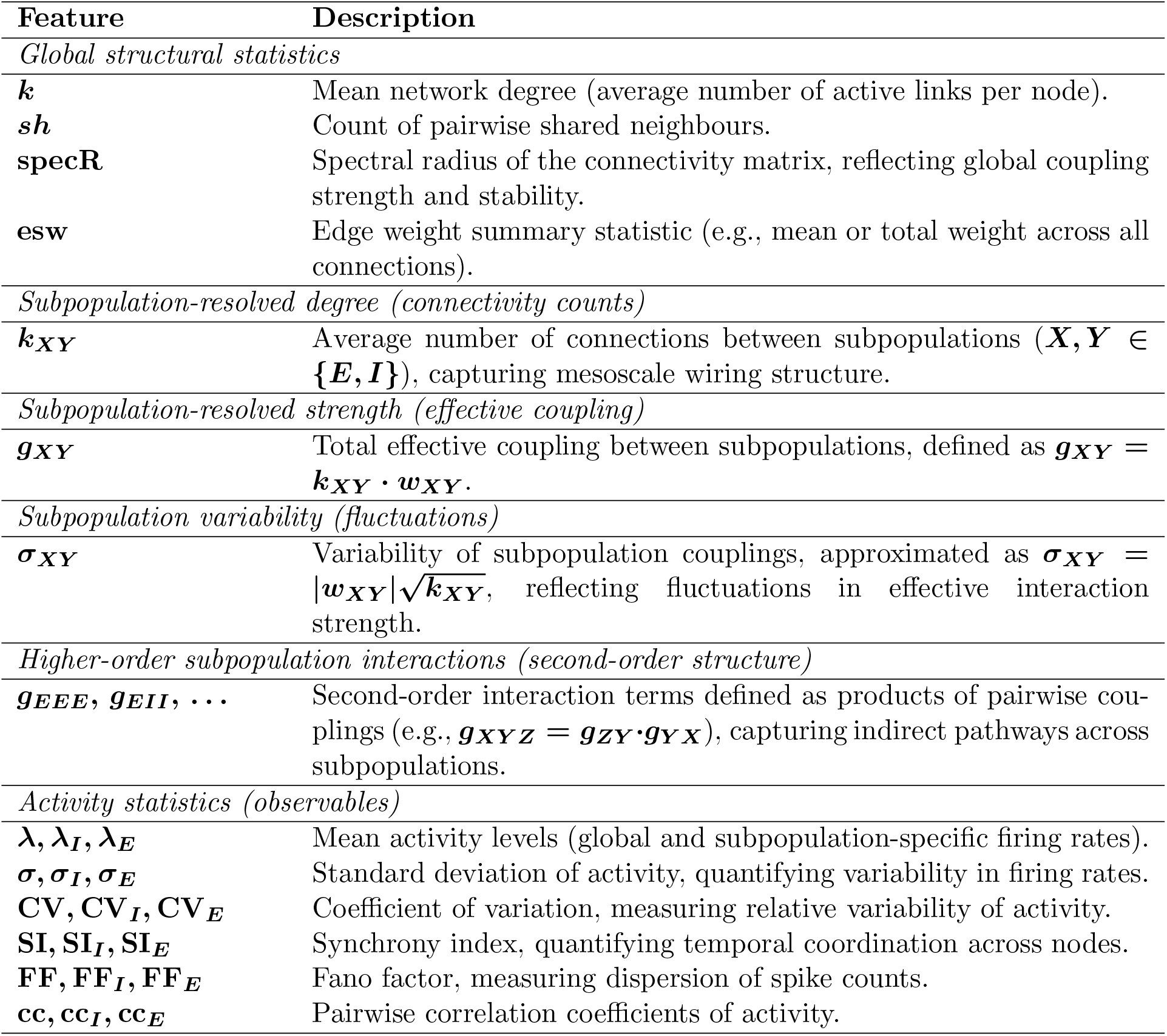
Summary of structural and functional features used to construct correlation matrices. Features are grouped by their level of description, from global statistics to higher-order subpopulation interactions. Subscripts ***E*** and ***I*** denote excitatory and inhibitory populations, respectively.

#### Factorial design and dataset composition

Both dataset families shared the same factorial structure: weighting scheme, network class, realization, synaptic weight scale, degeneration mode (synaptic vs neuronal), pruning strategy, and degeneration stage. For each family, the two weighting schemes were stored as separate arrays, so the relevant multiplicative factors were: 2 weighting-scheme arrays, 6 network classes, 10 realizations, 10 synaptic weight scales, 2 degeneration modes, 5 pruning strategies, and 10 degeneration stages.

For either dataset family, the number of intact weighted baseline networks was

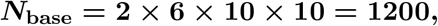

and for each baseline we generated

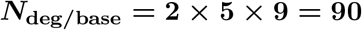

degeneration-derived networks, yielding a total of

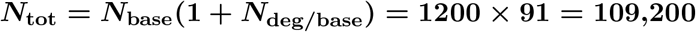

unique network states.

Because the tuned and untuned families had the same dimensions in the stored arrays, the tuned family contributed the same number of unique states as the untuned family:

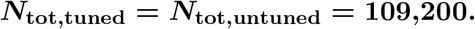

Across both dataset families, the total number of unique network states analyzed in this study was

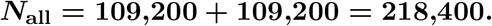

### Neuron and synapse model

#### Neuron model

Neuronal dynamics were modeled using a current-based leaky integrate-and-fire (LIF) neuron. The subthreshold membrane potential obeyed

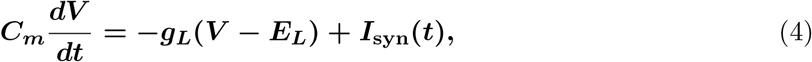

where ***C***_***m***_ is the membrane capacitance, ***g***_***L***_ **= *C***_***m***_***/τ***_***m***_ the leak conductance, ***τ***_***m***_ the membrane time constant, and ***E***_***L***_ the resting potential. When ***V* (*t*)** reached the threshold ***V***_**th**_, a spike was emitted, after which the membrane potential was reset to ***V***_**reset**_ and held refractory for a duration ***t***_**ref**_. Supplementary Table 2 lists the main parameters used in the study.

#### Synaptic model

Synaptic interactions were modeled as current-based alpha synapses. Each presynaptic spike from neuron ***j*** to neuron ***i*** generated a postsynaptic current

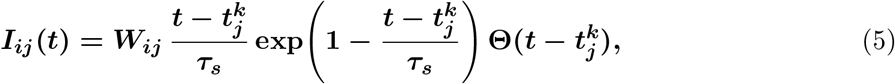

where ***W***_***ij***_ is the signed synaptic weight, ***τ***_***s***_ the synaptic time constant, 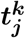 the ***k***-th spike time of neuron ***j***, and **Θ(·)** the Heaviside function. The total synaptic input current to neuron ***i*** is

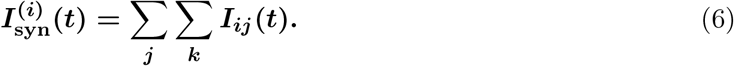

### Simulation protocol and activity measures

Each network instance was simulated for **11** s of biological time with temporal resolution **Δ*t* =0.1** ms. An initial transient of ***T***_**trans**_ **= 1** s was discarded, and all analyses were performed on the remaining ***T* = 10** s window.

Neuronal activity was quantified using complementary measures capturing firing rate, firing-rate variability, spike-time irregularity, population variability, pairwise correlations, and subthreshold synchronization.

#### Firing rate

The mean firing rate was

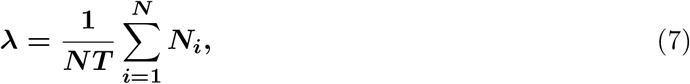

where ***N***_***i***_ is the number of spikes emitted by neuron ***i*** during the analysis window of duration ***T***. Population variability of firing rates was quantified by the standard deviation ***σ***_***λ***_ across neurons.

#### Spike-time irregularity

Spike-time irregularity was quantified by the coefficient of variation (CV) of inter-spike intervals (ISIs). For neuron ***i***,

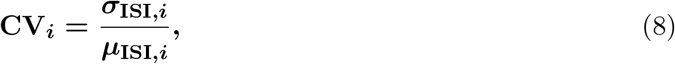

and population-level CV is the mean of **CV**_***i***_ across neurons with sufficient spike counts.

#### Population-level asynchrony

Population asynchrony was assessed using the Fano factor (FF) of the population spike-count histogram. Spikes were binned into windows of width **Δ*t***_***b***_ **= 1** ms and

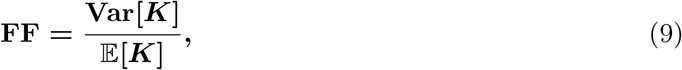

where ***K*** is the spike count per bin.

#### Pairwise spike-count correlations

Pairwise correlation coefficients (cc) were computed by binning spike trains into windows of width **Δ*t***_***b***_ **= 1** ms and evaluating Pearson correlation coefficients between pairs of binned spike-count time series. Reported values correspond to averages over all neuron pairs.

#### Synaptic current

Synaptic current (SynI) refers to the total synaptic input current received by a neuron, obtained as the sum of all incoming excitatory and inhibitory alpha-shaped synaptic currents during the analysis window.

#### Structural descriptors

Structural descriptors were computed directly from the pruned weighted adjacency ***W*** ^**(*i*)**^ at each degeneration stage.

#### Degree

In-degree and out-degree were computed from the binary adjacency ***A***^**(*i*)**^, and population-level statistics were obtained by averaging across neurons.

#### Shared presynaptic count

For neurons ***n***_***i***_ and ***n***_***j***_, the number of shared presynaptic neurons was

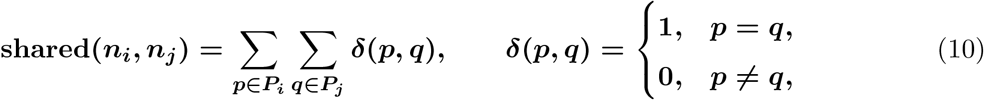

where ***P***_***i***_ and ***P***_***j***_ are the presynaptic sets of ***n***_***i***_ and ***n***_***j***_, respectively. The network-level shared-input measure was obtained by averaging **shared(*n***_***i***_, ***n***_***j***_) over all neuron pairs.

#### Effective synaptic weight (ESW)

The effective synaptic weight of neuron ***i*** was defined as the sum of absolute incoming synaptic weights,

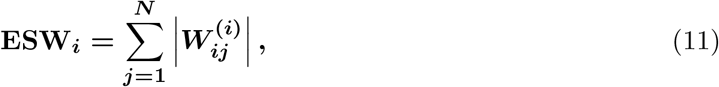

and the population-level ESW was obtained by averaging **ESW**_***i***_ across neurons.

#### Spectral radius

The spectral radius **specR** was defined as the largest absolute eigenvalue of the weighted connectivity matrix ***W*** ^**(*i*)**^.

### Structural feature representation and prediction scheme

To relate network structure to activity in a unified and interpretable way, we constructed a compact population-level structural representation based on subpopulation connectivity and population sizes. Each network was characterized by subpopulation-level connection probabilities ***P* = (*p***_***EE***_, ***p***_***EI***_, ***p***_***IE***_, ***p***_***II***_), synaptic weights ***W* = (*w***_***EE***_, ***w***_***EI***_, ***w***_***IE***_, ***w***_***II***_), and population sizes **(*N***_***E***_, ***N***_***I***_). As above, indices follow the convention ***w***_***XY***_ denotes ***Y* → *X*** (presynaptic ***Y***, post-synaptic ***X***), and the same convention is used for the derived subpopulation drives ***g***_***XY***_. Expected in-degrees were defined as

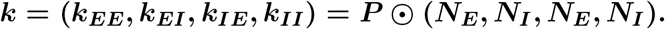

The mean synaptic drive was

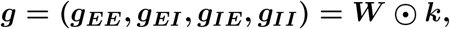

yielding four features. Fluctuation scales were approximated by

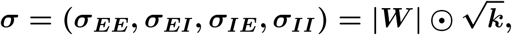

yielding four additional features. We approximated the variability of effective input from subpopulation ***Y*** to ***X*** using a mean-field scaling, 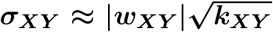, corresponding to independent unit-variance contributions from ***k***_***XY***_ inputs.

To capture nonlinear recurrent effects using two-step subpopulation interactions, we included eight second-order terms defined as products of subpopulation drives along length-2 population paths. Concretely, for a path ***X* → *Y* → *Z*** we defined

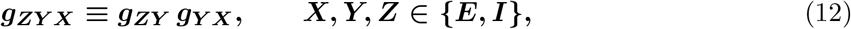

where ***g***_***YX***_ summarizes coupling from ***X* → *Y*** and ***g***_***ZY***_ summarizes coupling from ***Y* → *Z***, so that the product represents the composite path ***X* → *Y* → *Z***. For example,

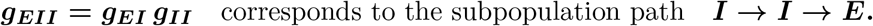

The remaining interaction terms were defined analogously for the other population triplets. Collectively, the **4** mean-drive features, **4** fluctuation-scale features, and **8** interaction features form a 16-dimensional structural representation.

All activity measures were predicted from the same 16-dimensional feature space using ridge regression with standardized features and an unpenalized intercept. Prediction performance was evaluated using repeated random train–test splits (25% test data) and leave-one-size-out (LOSO) cross-validation. Model accuracy was quantified using the coefficient of determination ***R***^**2**^.

### Reproducibility and validation

We enforced deterministic seeds for randomized steps (network generation, rewiring, and pruning). Validation checks included edge–matrix round-trip consistency, correctness of maximum-matching layering, monotone decrease in adjacency density under synaptic pruning, and monotone decrease in surviving node count under neuronal pruning. Automated regression tests ensured reproducibility.

## Data and Code Availability

All network generation, degeneration, simulation, and analysis code used in this study is available at https://github.com/absima/netDegenerate. The repository includes scripts for network construction, pruning strategies, weighting schemes, and computation of structural and dynamical measures. Parameters required to reproduce the simulations are provided in the documentation.

## Acknowledgments

This work was supported by the PEPR Santé Numérique program (France 2030), project “Brain Health Trajectories (BHT)”, implemented by the Agence Nationale de la Recherche (ANR) under grant number ANR-22-PESN-0012-BHT. We wish to thank Itamar D. Landau and collaborators for sharing the experimental microcircuit data.

## Supplementary Information

**Figure S1.**
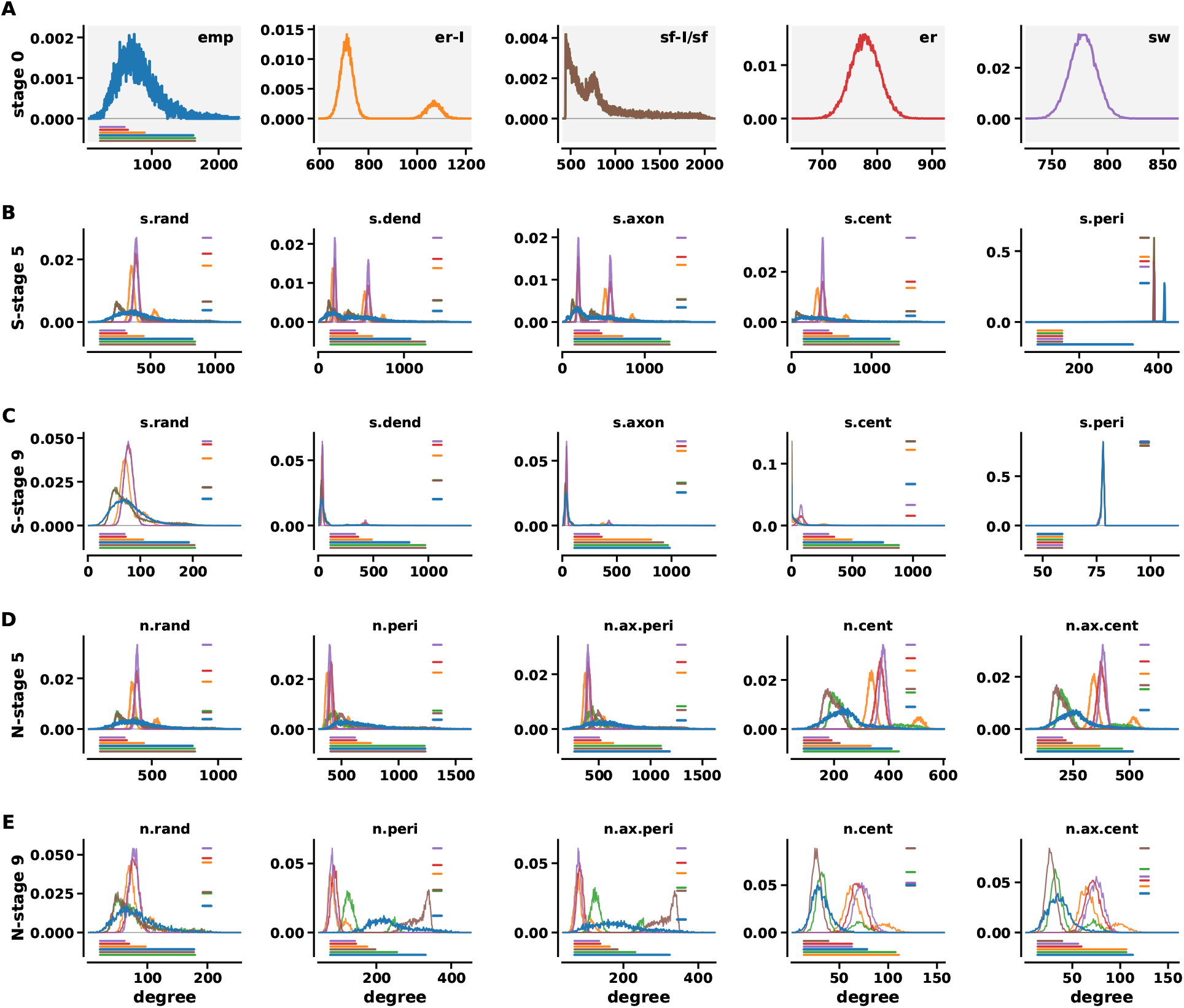
Degeneration-induced changes in degree distributions across network classes. Degree distributions of excitatory and inhibitory populations are shown for all network classes under different degeneration regimes. **(A)** Baseline (stage 0) degree distributions prior to degeneration, corresponding to the joint excitatory and inhibitory population profiles shown in Figure 1B. **(B–C)** Synaptic pruning at intermediate (stage 5) and advanced (stage 9) degeneration for all pruning strategies (`s.rand, s.dend, s.axon, s.cent, s.peri`). **(D–E)** Neuronal pruning at stage 5 and stage 9 for all strategies (`n.rand, n.peri, n.ax.peri, n.cent, n.ax.cent`). Across classes, degeneration progressively reshapes degree distributions in a strategy-dependent manner. Synaptic pruning primarily reduces connectivity while preserving the overall population size, leading to gradual shifts and, in some cases, sharpening of distributions depending on pruning priority. In contrast, neuronal pruning induces more pronounced structural reorganization by removing nodes, resulting in stronger distortions and truncation of degree distributions. Hub-targeted (central) strategies produce rapid collapse of high-degree tails, whereas peripheral strategies preferentially eliminate low-degree nodes, effectively narrowing or shifting the distribution. These effects highlight how different degeneration mechanisms redistribute connectivity rather than uniformly reducing it.

**Figure S2.**
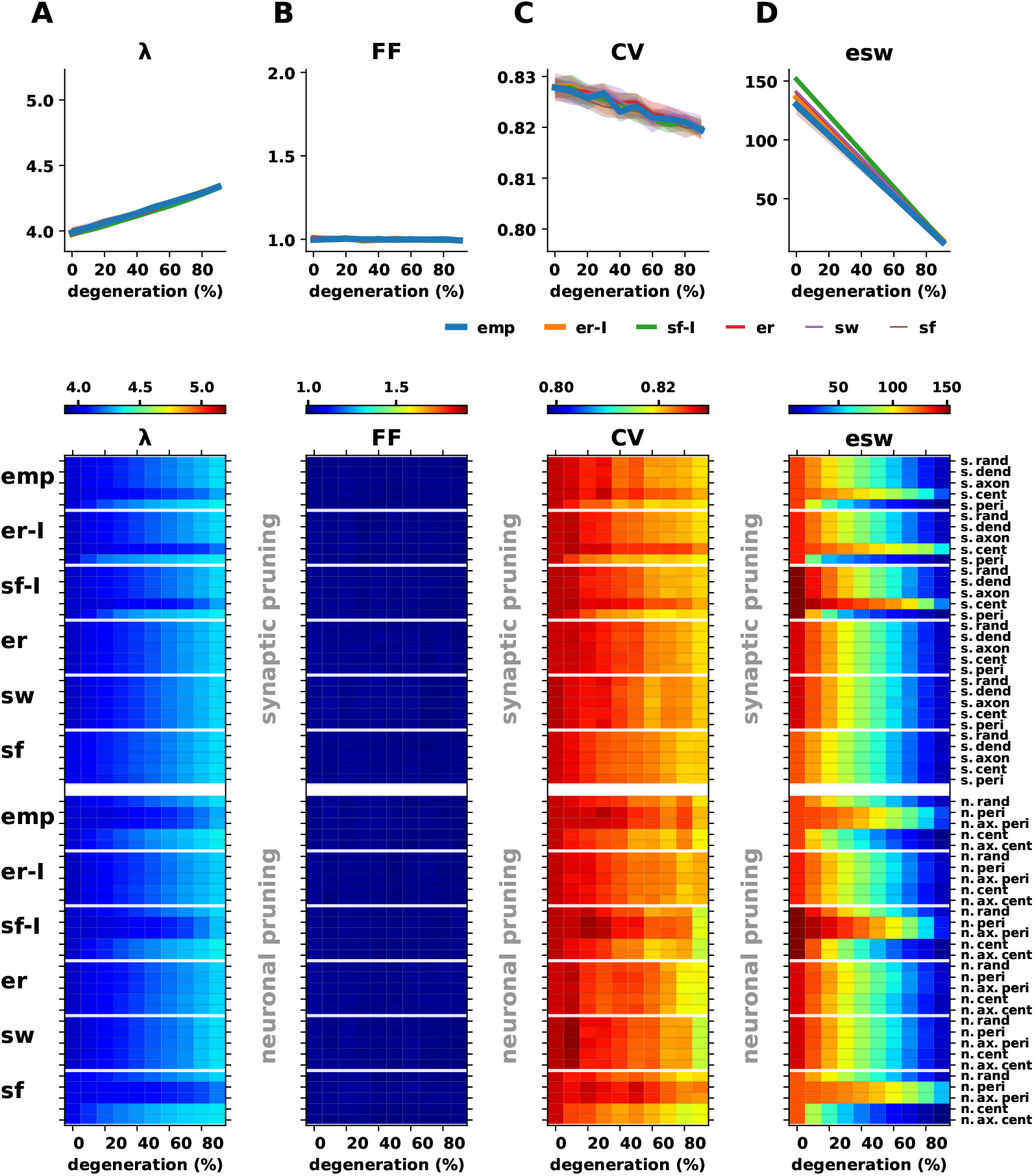
Strong inhibition reduces architecture-dependent differences in degeneration responses. Layout as in Fig. 2, with identical color scales used to enable direct cross-comparison between the two inhibition regimes. Columns correspond to mean firing rate (A), population Fano factor (B), ISI coefficient of variation (C), and absolute effective synaptic weight (D). Top row shows degeneration trajectories for random synaptic pruning, and the heatmaps summarize responses across pruning strategies for the tuned baseline condition (4 Hz), with synaptic pruning in the upper panel and neuronal pruning in the lower panel. In this stronger inhibition regime, inhibitory synapses dominate network dynamics, and degeneration responses become more similar across all six classes. The architecture-dependent differences observed in Fig. 2 are therefore substantially reduced.

**Figure S3.**
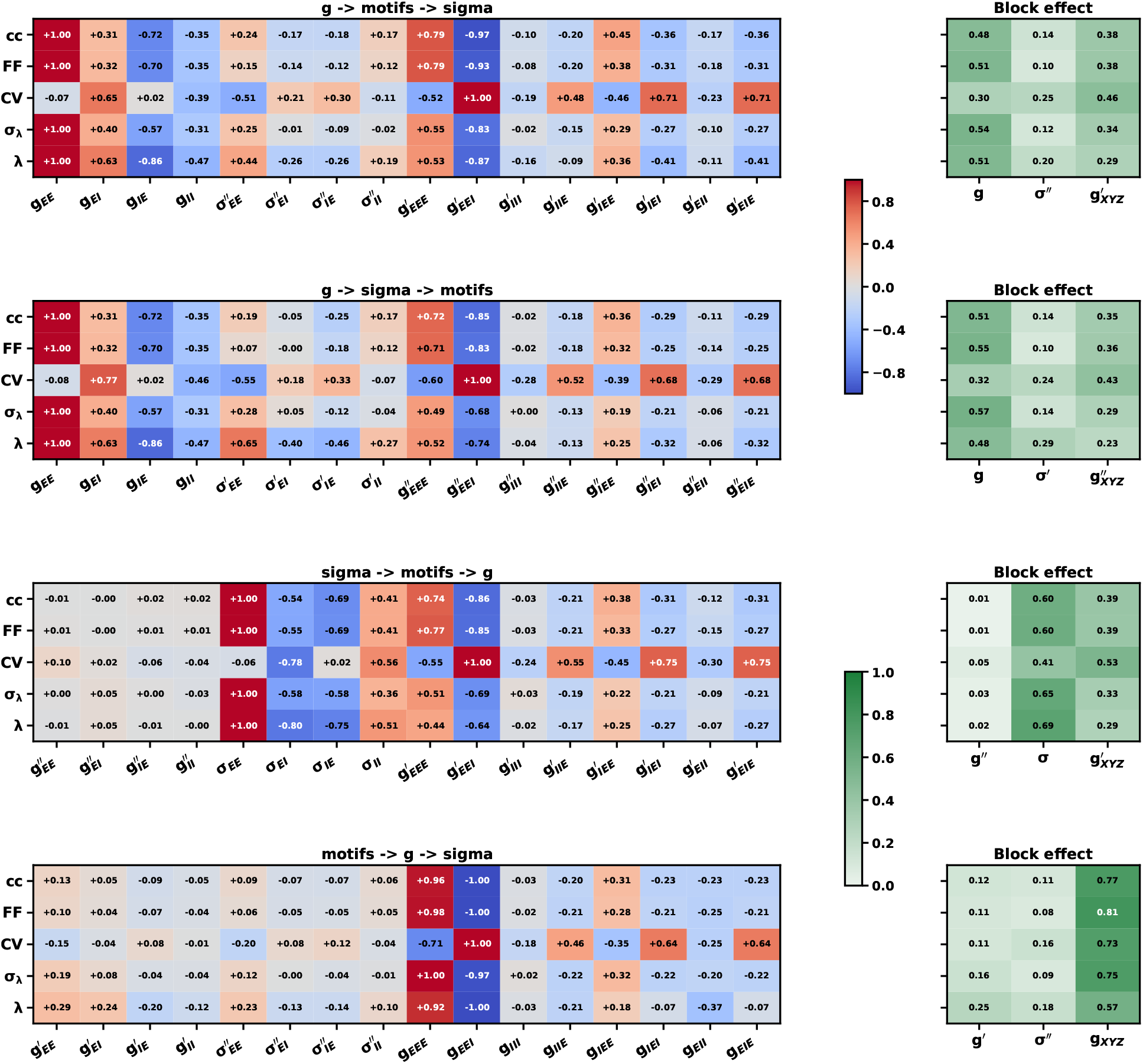
Order-dependent orthogonalized contributions of subpopulation feature families. Rows correspond to activity measures (pairwise spike-count correlation ***cc***, population Fano factor ***F F***, spike-time irregularity ***CV***, firing-rate variability ***σ***_***λ***_, and mean firing rate ***λ***). Each large heatmap shows one sequential decomposition of the three weight-aware feature families: subpopulation mean drives ***g***_***XY***_, fluctuation scales ***σ***_***XY***_, and second-order interaction terms ***g***_***XYZ***_. The titles indicate the order in which feature families were introduced; later families were orthogonalized with respect to the span of earlier ones before their contributions were evaluated. Entries in the main heatmaps show the resulting signed feature contributions for the corresponding ordering. The small panel to the right of each heatmap summarizes the aggregate contribution of each feature family as a **block effect**. Across orderings, mean drives and fluctuation scales capture strongly overlapping predictive information, whereas interaction terms retain a smaller but more distinct contribution once lower-order features have been accounted for.

**Figure S4.**
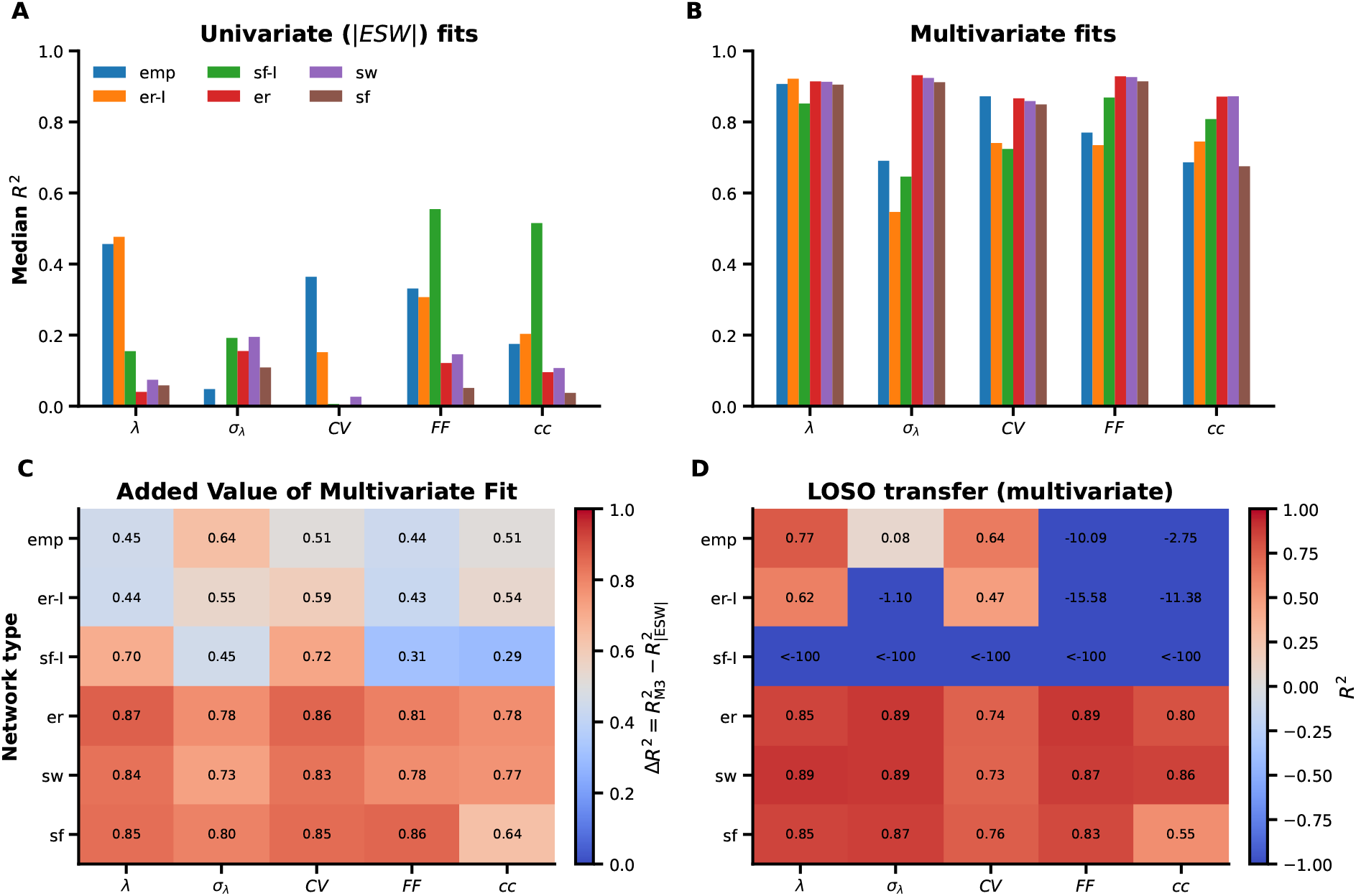
Univariate and multivariate structure–function fits capture complementary aspects of degeneration dynamics. **(A)** Median within-network prediction performance (***R***^**2**^) of the univariate |**ESW**| model for mean firing rate (***λ***), firing-rate variability (***σ***_***λ***_), spike-time irregularity (***CV***), population Fano factor (***F F***), and pairwise spike-count correlation (***cc***), shown separately for each network class. **(B)** The corresponding within-network performance of the multivariate ridge model based on the 16-dimensional weight-aware structural feature set. **(C)** Within-network gain of the multivariate description relative to the univariate one, quantified as 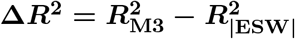 for each class and observable. Positive values throughout show that the multivariate representation consistently improves prediction even when evaluation is restricted to individual classes. **(D)** Leave-one-network-out transfer performance of the multivariate model, where each row denotes the test class held out during training and each entry reports the resulting test-set ***R***^**2**^. Transfer remained strong for `er, sw`, and `sf`, but degraded substantially for sf-I, indicating that although the multivariate representation captured much more variance than |**ESW**| alone, its generalization across architectures remained class-dependent. Taken together, these comparisons show that |**ESW**| retains informative class-level predictive value, but multivariate weight-aware descriptors provide a substantially richer account of degeneration-induced activity changes.

## References

[1] S. T. DeKosky and S. W. Scheff. Synapse loss in frontal cortex biopsies in alzheimer’s disease: correlation with cognitive severity. Annals of Neurology, 27(5):457–464, 1990.

[2] R. D. Terry, E. Masliah, D. P. Salmon, and et al. Physical basis of cognitive alterations in alzheimer’s disease: Synapse loss is the major correlate of cognitive impairment. Annals of Neurology, 30(4):572–580, 1991.

[3] Dennis J. Selkoe. Alzheimer’s disease is a synaptic failure. Science, 298(5594):789–791, 2002.

[4] M. Day, Z. Wang, J. Ding, and et al. Selective elimination of glutamatergic synapses on striatopallidal neurons in parkinson disease models. Nature Neuroscience, 9(2):251–259, 2006.

[5] A. J. Milnerwood and L. A. Raymond. Early synaptic pathophysiology in neurodegeneration: insights from huntington’s disease. Trends in Neurosciences, 33(11):513–523, 2010.

[6] W. Shen and et al. Striatal synaptic adaptations in parkinson’s disease: mechanisms and functional implications. Trends in Neurosciences, 45(6):433–447, 2022.

[7] Nimeshan Geevasinga, Pubudu Menon, and et al. Pathophysiological and diagnostic implications of cortical hyperexcitability in als. Clinical Neurophysiology, 127(6):2547–2560, 2016.

[8] Ed Bullmore and Olaf Sporns. The economy of brain network organization. Nature Reviews Neuroscience, 13(5):336, 2012.

[9] Mark Newman. Networks. Oxford university press, 2018.

[10] Eve Marder and Jean-Marc Goaillard. Variability, compensation and homeostasis in neuron and network function. Nature Reviews Neuroscience, 7(7):563–574, 2006.

[11] Friedemann Zenke, Wulfram Gerstner, and Surya Ganguli. The temporal paradox of hebbian learning and homeostatic plasticity. Current opinion in neurobiology, 43:166–176, 2017.

[12] Carl Van Vreeswijk and Haim Sompolinsky. Chaos in neuronal networks with balanced excitatory and inhibitory activity. Science, 274(5293):1724–1726, 1996.

[13] Nicolas Brunel. Dynamics of sparsely connected networks of excitatory and inhibitory spiking neurons. Journal of computational neuroscience, 8(3):183–208, 2000.

[14] Arvind Kumar, Sven Schrader, Ad Aertsen, and Stefan Rotter. The high-conductance state of cortical networks. Neural computation, 20(1):1–43, 2008.

[15] Itamar D Landau, Robert Egger, Vincent J Dercksen, Marcel Oberlaender, and Haim Sompolinsky. The impact of structural heterogeneity on excitation-inhibition balance in cortical networks. Neuron, 92(5):1106–1121, 2016.

[16] Tim Fieblinger, Steven M Graves, Luke E Sebel, Cristina Alcacer, Joshua L Plotkin, Tracy S Gertler, C Savio Chan, Myriam Heiman, Paul Greengard, M Angela Cenci, et al. Cell type-specific plasticity of striatal projection neurons in parkinsonism and l-dopa-induced dyskinesia. Nature communications, 5(1):5316, 2014.

[17] A. Mecca and et al. Synaptic density imaging in alzheimer’s disease: recent advances with sv2a pet. Alzheimer’s Research & Therapy, 2025. In press; update details once available.

[18] Pubudu Menon and et al. Cortical hyperexcitability and disease spread in amyotrophic lateral sclerosis: a longitudinal study. European Journal of Neurology, 24(12):1535–1543, 2017.

[19] J. Scekic-Zahirovic and et al. Cortical hyperexcitability in amyotrophic lateral sclerosis: human and mouse evidence. Science Translational Medicine, 16(838):adg3665, 2024.

[20] Rosa M. Villalba and Yoland Smith. Loss and remodeling of striatal dendritic spines in parkinson’s disease: from mechanisms to clinical implications. CNS Neuroscience & Therapeutics, 23(2):143–151, 2017.

[21] D. K. Wilton and et al. Complement-dependent elimination of corticostriatal synapses in huntington’s disease models. Nature Medicine, 29(5):1125–1135, 2023.

[22] Tara L. Spires-Jones and Bradley T. Hyman. The intersection of amyloid beta and tau at synapses in alzheimer’s disease. Neuron, 82(4):756–771, 2014.

[23] Soyon Hong, Violeta F. Beja-Glasser, Brianna M. Nfonoyim, Ana Frouin, Shan Li, Sruthi Ramakrishnan, Katherine M. Merry, Qiao Shi, Ann Rosenthal, Ben A. Barres, Cynthia A. Lemere, Dennis J. Selkoe, and Beth Stevens. Complement and microglia mediate early synapse loss in alzheimer mouse models. Science, 352(6286):712–716, 2016.

[24] Borislav Dejanovic, Melanie A. Huntley, Anne De Maziere, William J. Meilandt, Tony Wu, Kritika Srinivasan, Zora Modrusan Jiang, Vishal Gandham, Bruce A. Friedman, H. Ngu, Oded Foreman, Richard A. D. Carano, Betty Chih, Judith Klumperman, Corey Bakalarski, Jesse E. Hanson, Morgan Sheng, and Jesse E. Hanson. Changes in the synaptic proteome in tauopathy and rescue of tau-induced synapse loss by c1q antibodies. Neuron, 100(6):1322–1336, 2018.

[25] Alix de Calignon, Marcos Polydoro, Marc Suárez-Calvet, Caroline William, David H. Adamowicz, Kathy J. Kopeikina, Rose Pitstick, Naruhiko Sahara, Karen H. Ashe, George A. Carlson, Tara L. Spires-Jones, and Bradley T. Hyman. Propagation of tau pathology in a model of early alzheimer’s disease. Neuron, 73(4):685–697, 2012.

[26] Marc Colom-Cadena, Claire Davies, Sara Sirisi, Ji-Eun Lee, Elif M. Simzer, Makis Tzioras, Marta Querol-Vilaseca, et al. Synaptic oligomeric tau in alzheimer’s disease–a potential culprit in the spread of tau pathology through the brain. Neuron, 111(14):2170–2183, 2023.

[27] Han Fu, John Hardy, and Karen E. Duff. Selective vulnerability in neurodegenerative diseases. Nature Neuroscience, 21(10):1350–1358, 2018.

[28] Smita Saxena and Pico Caroni. Selective neuronal vulnerability in neurodegenerative diseases: from stressor thresholds to degeneration. Neuron, 71(1):35–48, 2011.

[29] Hirofumi Ozeki, Ian M. Finn, Evan S. Schaffer, Kenneth D. Miller, and David Ferster. Inhibitory stabilization of the cortical network underlies visual surround suppression. Neuron, 62(4):578–592, 2009.

[30] Yashar Ahmadian, Daniel B. Rubin, and Kenneth D. Miller. Analysis of the stabilized supra-linear network. Neural Computation, 25(8):1994–2037, 2013.

[31] Daniel B. Rubin, Stephen D. Van Hooser, and Kenneth D. Miller. The stabilized supralinear network: a unifying circuit motif underlying multi-input integration in sensory cortex. Neuron, 85(2):402–417, 2015.

[32] William W. Seeley, Richard K. Crawford, Juan Zhou, Bruce L. Miller, and Michael D. Greicius. Neurodegenerative diseases target large-scale human brain networks. Neuron, 62(1):42–52, 2009.

[33] Cornelis J. Stam. Modern network science of neurological disorders. Nature Reviews Neuroscience, 15(10):683–695, 2014.

[34] Yang-Yu Liu, Jean-Jacques Slotine, and Albert-László Barabási. Controllability of complex networks. nature, 473(7346):167–173, 2011.

[35] Simachew Abebe Mengiste, Ad Aertsen, and Arvind Kumar. Effect of edge pruning on structural controllability and observability of complex networks. Scientific reports, 5(1):18145, 2015.

